# Generating long deletions across the genome with pooled paired prime editing screens

**DOI:** 10.1101/2025.11.03.686307

**Authors:** Juliane Weller, Gareth Girling, Elin Madli Peets, Jonas Koeppel, Seri Kitada, Pierre Murat, Fabio Giuseppe Liberante, Yousra Belattar, Sho Iwama, Tom Ellis, Leopold Parts

## Abstract

Engineered deletions are a powerful probe for studying genome architecture, function, and regulation. Yet, the lack of effective methods to create them in large numbers and at multi-kilobase scale has rendered interrogating all the gigabases of human DNA elusive. Prime editing based approaches are promising contenders to fill this gap, but have not been adopted for large-scale pooled screening so far. Here, we performed the first high-throughput paired prime editing deletion screens. To establish the technology, we generated 3,802 deletions of up to 1.2 Mb in a reporter locus in two human cell lines, and identified key determinants of paired prime editing efficiency to be deletion length, predicted pegRNA efficacy, and contact frequency. We used these learnings to generate therapeutically important deletions at rates that are expected to give clinical benefits for retinitis pigmentosa and Duchenne muscular dystrophy. Combining several technological advances, we then performed the first pooled paired prime editing screen to delete over 10,471 sequences of median length 1.1 kb that target 149 genes across the genome. Our screens were high quality, recovering 75% of positive controls. We discovered 462 non-coding regions with fitness effects, and validated deletions with impact as well as tolerated ones in regions with functional annotations. Our screens are the largest pooled characterization of precise deletions to date, demonstrating that prime deletion screens can perturb the genome at scale, and highlighting that most non-coding DNA around essential genes is dispensable for cell growth in one condition.

## Introduction

Understanding the function of all of the human genome requires tools that can interrogate large regions of DNA at scale. While protein-coding regions constitute less than 2% of the genome, up to 10% of the remaining non-coding DNA is estimated to be involved in regulatory and structural roles (Rands et al. 2014; ENCODE Project, Leja, and Birney 2012). This includes enhancers, promoters, non-coding RNAs, and chromatin-associated elements, many of which act during development (Jindal and Farley 2021) and have been implicated in disease (Stankiewicz and Lupski 2010). Despite these observations, direct validation of function of non-coding elements remains limited, in part because we lack high-throughput tools to precisely and scalably manipulate large genomic segments.

Targeted deletions are an effective strategy to study non-coding regions, as they enable the removal of entire elements or large stretches of DNA in a single step. Historically, laborious engineering with recombinases was used to remove single large DNA segments (Koeppel et al. 2024). More recently, the CRISPR-Cas9 system has been applied efficiently in pooled screens to induce deletions through paired guide RNAs that flank the desired region (Gasperini et al. 2017; Diao et al. 2017; Li et al. 2024; Aregger, Xing, and Gonatopoulos-Pournatzis 2021). While this method is scalable and conceptually simple, it relies on the stochastic repair of double-strand breaks, which is far from guaranteed to delete the flanked sequence rather than vandalize the targeted sites (Gasperini et al. 2017). The DNA breaks also introduce toxicity (Jackson 2002), off-target effects (Doench et al. 2016; X.-H. Zhang et al. 2015), and unpredictable chromosomal abnormalities (Kosicki, Tomberg, and Bradley 2018), which complicate the interpretation of the observed signal. The related CRISPR interference and epigenetic silencing approaches are useful for mapping regulatory regions (Fulco et al. 2016), but edit broadly around a locus (Usluer et al. 2023), and do not offer the range and precision to resolve large or overlapping functional elements.

Prime editing has emerged as a powerful alternative editing tool, offering programmable genome manipulation without double-strand breaks or donor DNA templates (Anzalone et al. 2019). By fusing a reverse transcriptase to a nicking Cas9, and guiding the complex with engineered prime editing guide RNAs (pegRNAs), prime editing enables precise base changes, insertions, or deletions. Paired prime editing extends this approach by using two pegRNAs to target opposite strands, generating flaps that replace intervening sequences (Anzalone et al. 2022; Choi et al. 2022; Kweon et al. 2023). This method enables clean deletions and, if desired, simultaneous insertions of sequences such as recombinase sites (Pandey et al. 2024). While paired prime editing has been demonstrated in small-scale and locus-specific contexts, its efficiency in pooled screens has been reported to be low (Cirincione et al. 2025; Herger et al. 2025), and its potential for deletion screens has not yet been realized.

A major barrier to scaling prime deletion screens is the variable efficacy of pegRNAs that specify the target and the outcome. This in turn is due to limited understanding of the factors that govern the editing efficiency, especially for large deletions. Previous studies have characterized pegRNA design parameters and chromatin context in single-edit contexts (Mathis et al. 2023, 2024; Koeppel et al. 2023), but the interactions between paired guides and the impact on the deleted genomic region remain poorly understood. Variables such as deletion length, local 3D contact frequency, and pegRNA compatibility likely influence outcomes, yet have not been systematically investigated. Furthermore, no prior study has demonstrated that paired prime editing can achieve megabase-scale deletions in a pooled format or be applied to disease-relevant loci at therapeutic scales.

Here, we perform the first high-throughput deletion screens using paired prime editing. We begin by establishing the capacity of the method in a model system, generating over 3,800 deletions of up to 1.2 Mb in two human cell lines. From this screen, we uncover key determinants of prime deletion efficiency, including deletion length, pegRNA efficacy, and 3D contact frequency, and use this information to engineer clinically meaningful deletions in models of Duchenne muscular dystrophy and retinitis pigmentosa. Building on these findings, we perform a genome-scale screen of over 10,000 deletions across 149 genes, achieving ∼72% recall of positive controls and identifying hundreds of non-genic and non-coding regions with fitness consequences. Our work demonstrates the scalability and precision of paired prime editing and provides a way forward for large-scale functional interrogation of the entirety of the human genome.

## Results

We set out to delete large stretches of the human genome using paired prime editing. To do so, we first measured the efficacy of generating deletions of different sizes and qualities with the aid of a reporter screen. We used the HAP1 and HEK293T cell lines with the mScarlet red fluorescent protein integrated into the CLYBL locus, chose three sites as the deletion start points with an anchor epegRNA, and designed a pooled epegRNA library to define deletion end points, aiming to generate 3,802 diverse deletions ranging from 45 bp to 1.2 Mb (Figure 1b, Figure S1, Table S1). We screened pooled libraries against each anchor across both cell lines by introducing epegRNA pairs, inducing editing, sorting for reporter fluorescence, and sequencing DNA to measure the abundance of each epegRNA after reporter disruption (Figure 1c). To quantify the deletion generation efficacy, we calculated normalized log_2_ fold changes (LFCs, Table S1) between the sorted low and high reporter expression pools. The LFCs were reproducible (Figure S2), with median replicate Pearson’s correlation of 0.57 (range: 0.38-0.79), and increasingly high correlations when combining data from replicates (median R=0.68 between same anchors in different cell lines, median R=0.72 between different anchors in same cell line; Figure 1d), and anchors (R=0.79 between cell lines when combining replicates; Figure 1e). For the remaining analyses, we use the average signal across all replicates, anchors, and cell lines.

**Figure 1.**
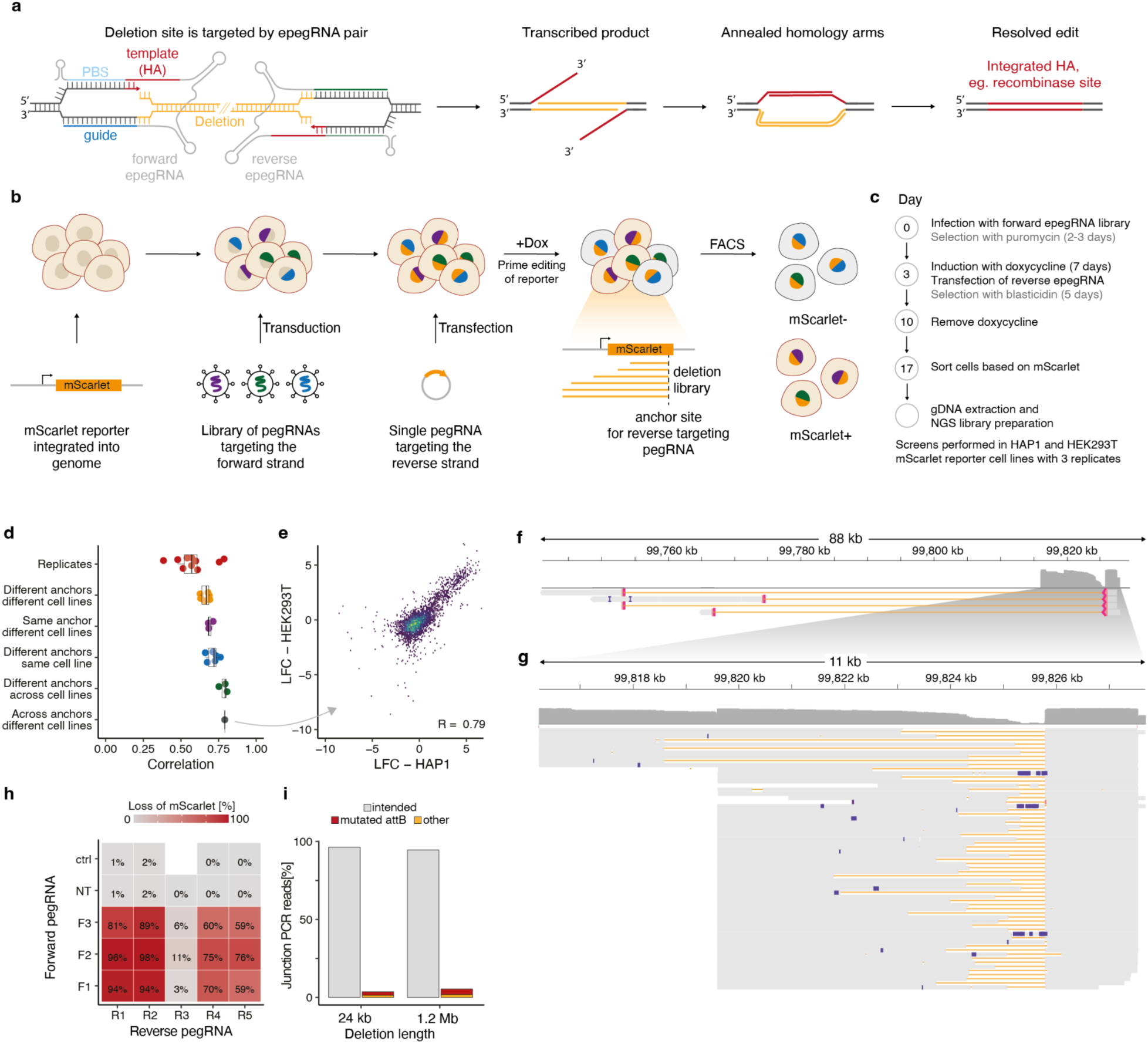
Screening and validating paired prime editing for generating reporter deletions at scale. **a**. Outline of paired prime editing for generating long deletions. **b**. Outline of screening approach for quantifying paired prime deletion efficacy. **c**. Timeline and selection regimes for the screen. **d**. Correlation (x-axis) of different types of samples (y-axis, colors) for individual comparisons (markers). Comparisons are all-against-all, replicates are combined for all but the first set of samples. Box: median and quartiles; whiskers: 1.5x interquartile mean. **e**. Average log_2_ fold-change between reporter positive and negative populations in HEK293T cells (x-axis) and HAP1 cells (y-axis) for individual deletions (markers). **f**. Coverage (grey, y-axis) along the engineered locus (x-axis) of nanopore sequencing reads. **g**. As (f), but focusing on the 11kb region around the cut site, highlighting individual reads (lines). Grey: well covered sequence; orange: deleted sequence; blue markers: called mutations. **h**. Fraction of reporter negative cells indicating deletion efficiency (color, numbers) for combinations of forward (y-axis) and reverse (x-axis) pegRNAs. Ctrl: no forward pegRNA; NT: non-targeting pegRNA. **i**. Percentage of observed reads (y-axis) with intended outcomes (grey), mutated attB sites (red), or other mutations (yellow) for different deletion lengths (x-axis).

The screen results suggested that we can successfully generate deletions, with on average 16% of cells losing mScarlet fluorescence. To validate that this loss was driven by deletions rather than silencing or small mutations, we performed targeted PCR-free long-read sequencing of the reporter-negative cell population (Figure S3). We obtained direct sequence support from spanning long reads for deletions up to 70kb (Figure 1f), with most of the successful deletions less than 3kb long (Figure 1g). To confirm the high editing rates, we designed four forward-strand-targeting epegRNAs and five reverse-strand-targeting epegRNAs that could be combined pairwise to generate deletions of up to 1.5 kb (Table S2). Four of the five reverse epegRNAs edited with average rate of 80% across the forward epegRNAs (range: 59-98%, Figure 1h). One reverse epegRNA had low efficiency across tested combinations (3-11%, mean=6.5%), and the efficacy of negative controls of no reverse or non-targeting epegRNAs was below 1.8%, highlighting that both epegRNAs must be active to generate deletions efficiently. We also checked the purity of two generated deletions of sizes 24kb and 1.2Mb (Figure S4). In 2.9% of reads, the attB site was mutated and 1.6% of reads had a mutated target site (Figure 1i). Overall, the deletions were precise, with 95.5% of reads carrying the intended edit, which is at least as accurate as typical prime editing purities of 90-95% for PE2 systems (Anzalone et al. 2019).

To dissect the determinants of efficacy for generating long deletions, we explored the impact of epegRNA design features (Figure 2a, Table S3). The size of the generated deletion had a strong effect on the outcome, with efficacy diminishing with increasing deletion length (Figure 2b). Across the three anchors that specify the deletion start points, the editing efficiencies were highest for deletions up to 1 kb, with moderate success for deletions of 1-5kb, some signal for 5-10kb sizes, and small but significant difference from the negative control group for up to 100 kb deletions (Tukey HSD, p < 0.05; Figure 2c). For deletions longer than 100 kb, the difference from the controls was small and no longer statistically significant, suggesting a limit to generating large deletions at detectable rates in a screening setting. Focusing on variation between the rates of deletions up to 10kb, higher editing efficiencies were associated with better PRIDICT2 and DeepCas9 scores as well as the GC content of the primer binding site for the pegRNAs (R=0.39, 0.32, and 0.37, respectively, Figure 2d-f). Overall, the most efficiently generated deletions were up to 3kb, and overlap with the promoter and coding sequence of the expressed mScarlet reporter, likely reflecting the combined bias of the reverse-transcribed homology arms more easily annealing across smaller distances, and ease of editing in open and actively transcribed genomic contexts.

**Figure 2.**
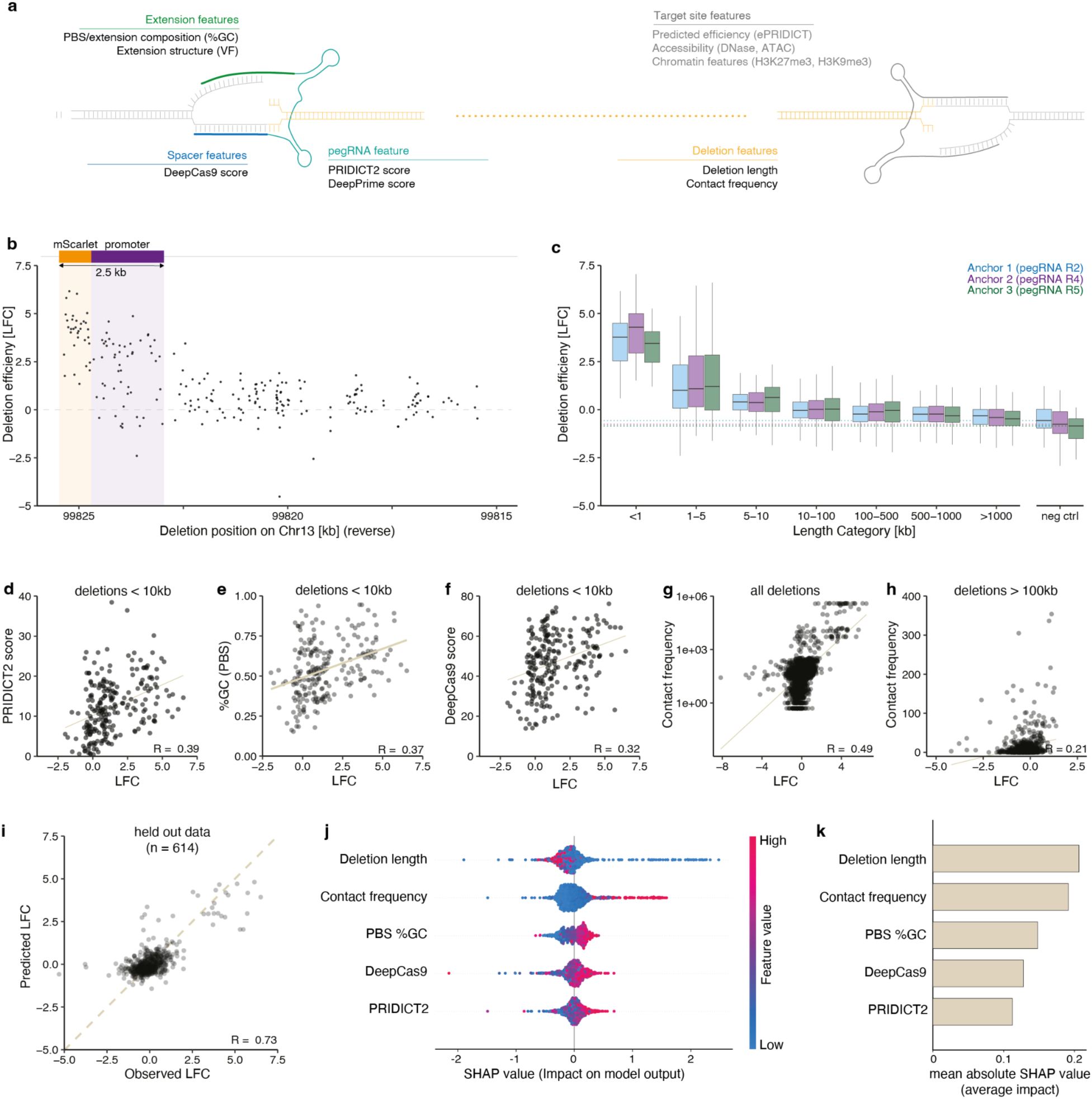
Determinants of paired prime editing efficiency for generating deletions. **a**. Features that may impact paired prime editing efficiency that were considered in the model. **b**. Log_2_ fold-change deletion signal (y-axis) observed along the reporter locus (x-axis) for R2 anchor pegRNA. Yellow shading: fluorescent reporter gene; purple shading: promoter region. **c**. Log_2_ fold-change deletion signal (y-axis) for pegRNAs stratified by their generated deletion length (x-axis) when paired with different anchor pegRNAs (colors). Box: median and quartiles; whiskers: 1.5 times the interquartile range. **d**. Log_2_ fold-change deletion signal (x-axis) contrasted against PRIDICT2 score (y-axis). Solid line: linear regression fit. R: Pearson’s correlation coefficient. **e**. As (d), but contrasting deletion signal against GC content in the primer binding site. **f**. As (d), but contrasting deletion signal against DeepCas9 score (y-axis). **g**. As (e), but contrasting deletion signal against MCC contact frequency. **h**. As (g), but focusing on deletions of at least 100kb. **i**. Predicted (y-axis) and observed (x-axis) log_2_ fold-change deletion signal for data points (markers) held out from model training. **j**. SHAP values (x-axis) for individual datapoints for features of the predictive model (y-axis). Color: sign of the impact on model output. **k.** Mean absolute SHAP values (x-axis) for predictive model features (y-axis).

To help distinguish between the impacts of 1-D distance along the linear genome, and 3-D distance of deletion endpoints in the nucleus for efficient deletion, we next quantified the contact frequency of the anchored deletion start site with the rest of the genome using micro-capture-C (MCC; Figure S5) (Hua et al. 2021). We confirmed that the MCC contact frequency is driven by distance to the probe, and that the target sites with high MCC scores also have the highest editing rates (Pearson’s R=0.49, Figure 2g). The positive correlation with deletion efficiency remains when subsetting the data to deletions over 100kb (Pearson’s R=0.21). While largely overlapping with deletion length, the MCC score can help distinguish candidate efficient deletions across larger distances.

Finally, we developed a model to predict deletion efficiencies. We fit gradient boosted regression models on training data (Methods), and achieved Pearson’s R=0.73 on held-out test data (Figure 2i). The SHAP values (Lundberg and Lee 2017) of the model highlight deletion length and contact frequency as the strongest contributing predictors, with spacer scores and pegRNA extension GC content as the remaining features with substantial impact (Figure 2j,k). Based on these findings, we recommend to target nearby or frequently contacting sites (with a 5kb upper distance limit) and to avoid epegRNAs with low predicted efficiency for generating deletions.

We next set out to use the uncovered design principles to generate individual clinically relevant ultra-long deletions. To do so, we first tested the efficiency of creating increasingly long deletions in isolation in HAP1 cells. We selected 24 high-performing forward-targeting epegRNAs from the screen, and attempted to delete 3kb to over 1Mb of the reporter with a fixed reverse epegRNA. The shortest deletion in this set had the highest editing efficiency (48%, Figure 3a), but also epegRNA pairs hundreds of kilobases apart resulted in substantial reporter loss. For example, a 252 kb deletion was generated in 6.5% of the cells, and even the three longest deletions exceeding 1 Mb were created at rates of 4.9%, 1.6%, and 3.5%. Meanwhile, the controls with single epegRNAs did not exceed 0.17% mScarlet loss. Thus, multi-kb deletions can be generated at high rates, and ultra-long deletions at lower rates.

**Figure 3.**
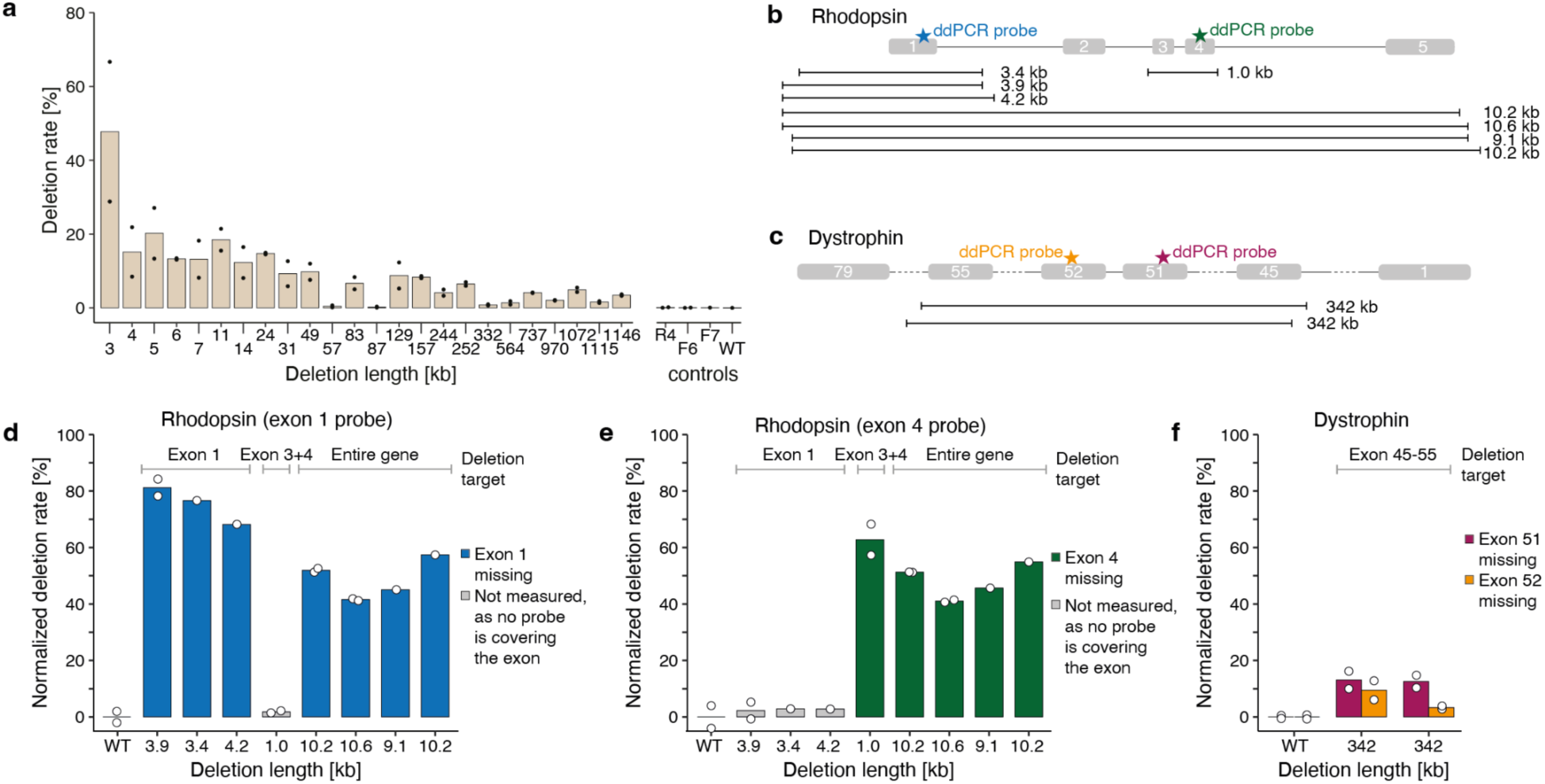
Long deletions can be created with paired prime editing at therapeutically relevant levels. **a**. Deletion rate as frequency of mScarlet reporter loss (y-axis) for different deletion lengths (x-axis). **b**. Outline of Rhodopsin gene deletion and measurement designs. **c**. Outline of Dystrophin gene deletion and measurement designs. **d**. Rhodopsin gene exon 1 deletion rate (y-axis) for pegRNA pairs targeting sets of exons (labels) to generate deletions of different sizes (x-axis). Markers: replicate values; WT: wild type negative control. **e**. As (c), but for exon 4 deletion rate. **f**. Deletion rate (y-axis) in the dystrophin gene (colors) for pegRNA pairs targeting different target sites (x-axis). Markers: replicate values; WT: wild type negative control.

There are several genetic diseases that could benefit therapeutically from generating large deletions. Retinitis pigmentosa, a progressive degenerative eye disease, is caused by dominant mutations in the *RHO* gene in 20–30% of cases. In principle, the mutated allele of the gene can be deleted, and then replaced via recombinase-mediated integration at the locus or via random integration (Humphries et al. 1997). Shorter deletions, such as those targeting individual exons, are more efficient. We attempted to delete both a single exon and the entire *RHO* gene using paired prime editing, and read out the efficacies by genotyping single cells with droplet digital PCR (Figure 3b). Targeting a single exon with three sets of epegRNAs, we achieved up to 84% efficiency for 3.4 – 4.2kb deletions, while deleting the entire gene in 9.1 – 10.6kb sections was successful 50% of the time on average (Figure 3d,e).

Duchenne muscular dystrophy (DMD) is a severe X-linked disorder characterized by progressive muscle weakness and degeneration. It is caused by mutations to the largest human gene dystrophin, with frameshifts in exons 45–55 making up ∼60% of pathogenic variants. The reading frame, and thus production of a partially functional protein, can be restored by deleting these exons (Aoki et al. 2012). We could engineer the required 340 kb deletion using paired prime editing at 5-13% efficiency (Figure 3e). This is a therapeutically meaningful level (Godfrey et al. 2015; Lu, Cirak, and Partridge 2014; Durbacz et al. 2025; Charleston et al. 2018), and offers an advantage over double-stranded break inducing methods of potentially reducing off-target effects, unwanted editing outcomes, and cytotoxicity.

After establishing the determinants of effective kilobase- to megabase-sized deletions, and demonstrating the ability to create therapeutic deletion alleles at clinically relevant frequencies, we turned to systematically interrogating the genome at scale. To do so, we implemented a pooled paired deletion screen to tile deletions that cover 10 kb around 149 selected genes of variable essentiality (Figure 4a; Table S4). We made use of the improved oligonucleotide synthesis abilities to design pairs of epegRNAs in a single construct of 433nt (Figure 4b), and the PE7 system that increases efficiency of prime editing (Yan et al. 2024) (Figure S6). We designed a library of 10,471 constructs with median intended deletion size 1kb according to the learnings above, and screened it for up to 28 days in the haploid HAP1 cells (Figure 4c). We calculated log_2_-fold changes (LFCs) of epegRNA pair frequencies between each timepoint and the initial one, with effective functional deletions expected to mostly drop out of the pool, and use these for analysis going forward.

**Figure 4.**
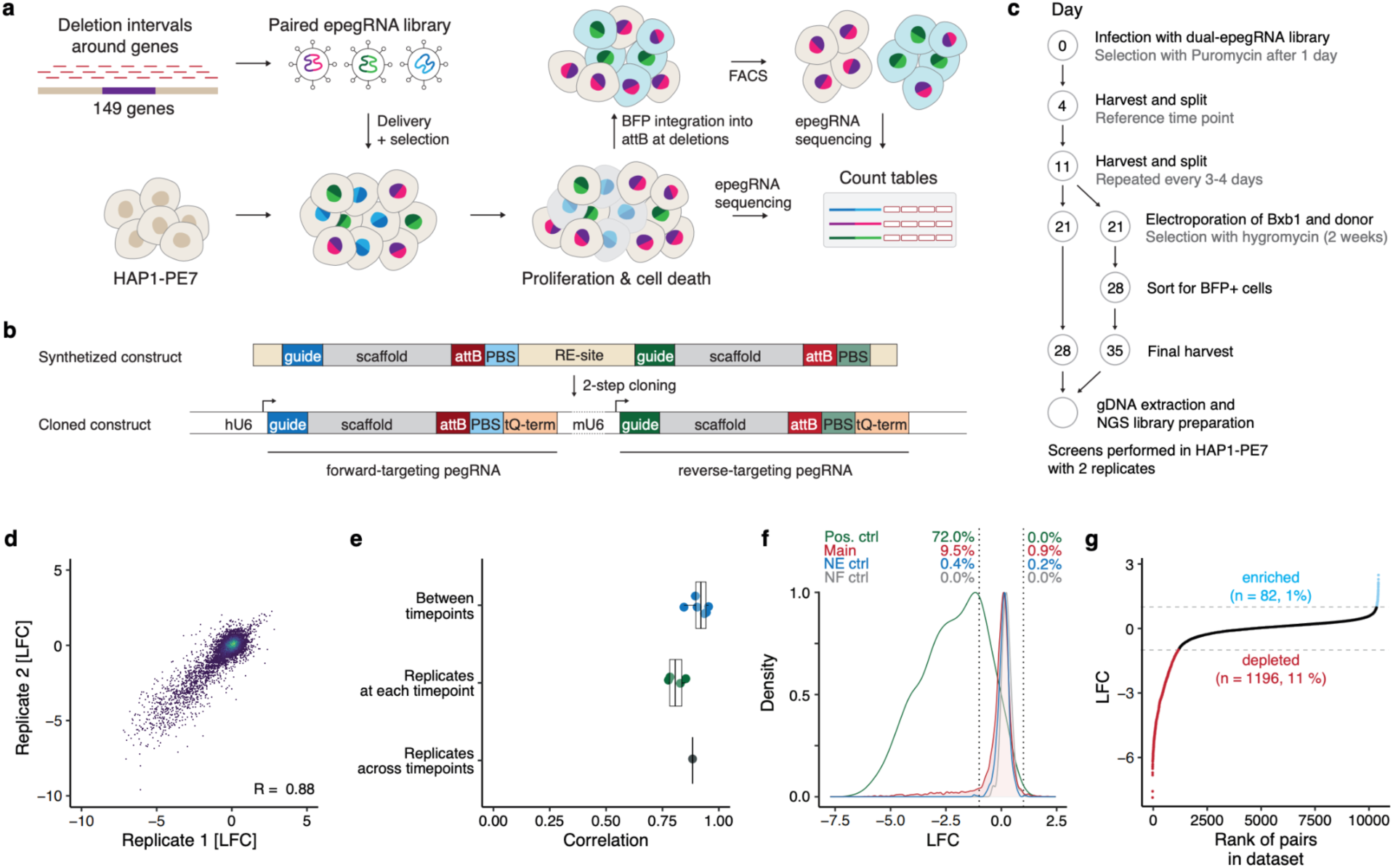
Pooled deletion screening of coding- and non-coding DNA around genes. **a**. Outline of pooled paired prime editing screen for phenotypic profile of long deletions genome-wide. **b**. Construct design and cloning strategy. **c**. Timeline and selection regimes for the screen. **d**. Phenotypic impact of individual deletions (LFC, markers) in two biological replicate experiments (x- and y-axis). R: Pearsons’s correlation coefficient. **e**. Correlation (Pearson’s R, x-axis) across combinations (y-axis) of timepoints and replicates. Markers: individual comparisons. Box: median and quartiles. **f.** Density (y-axis) of log2 fold changes (LFC, x-axis) for the non-essential control gene group (“NE”, blue), the non-functional control group (“NF”, gray), the positive control gene set with exonic deletions (“Pos. ctrl”, green), and the main screen (“Main”, red); dashed lines: +/- 1. Labels: fraction of hits above 1.0 (right) and below -1.0 (left) log2-fold change. **g**. Phenotypic impact (LFC, y-axis) of individual deletions (markers) sorted (x-axis) according to value. Red: depleted; blue: enriched; dashed lines: +/- 1.

We first confirmed that prime editing yields reproducible signal. The LFCs at timepoints along the screen correlated well across replicates (mean R=0.84, range 0.80-0.86, Figure 4d,e; Figure S7), as did the timepoints within a replicate screen (mean R=0.91, Figure 4e). As the LFCs were consistent, we merged the data first across timepoints (R=0.89 between replicates after merging), and then replicates for the remaining analyses. The final fitness estimates indicated that we have successfully mapped both deleterious and beneficial deletions, with 1,196 pegRNA pairs depleted (LFC < -1), 82 enriched (LFC > 1), and the majority neutral (Figure 4g). The non-targeting controls and non-essential gene targeting pegRNAs yielded hits at less than 1% rate (Figure 4f).

To quantify the screen’s sensitivity, we focused on deletions of essential gene coding regions. As expected, the majority of such deletions were depleted (LFC < -1; 72%, 724/1003, Figure 4f, 5a). The per-gene fraction of depleted exon deletions was 0.76 on average (Figure 5b,c), and correlated with gene essentiality estimated in a CRISPR knockout screen (Pearson’s R=0.79, Figure 5c). If a deletion was depleted, 68% of other deletions that overlapped it were also depleted (759/1,115 possible pairs concordant; Figure S8). Based on these signals, we estimate our recall of strongly essential regions to be ∼70% given the cutoffs used. The false negatives (deletions of essential gene exons not dropping out) could not be explained by their size, RNA expression from the gene, PRIDICT2 score, DeepCas9 score (maximum Pearson’s |R| with LFC of 0.18, Figure S8), or coding frame disruption (fraction of dropouts 0.69 vs 0.74 for frame preserving deletions vs not). Given our integration of principles of effective deletions into the library design that restricted the range of these parameters, we expected little impact of their residual variation on deletion efficiency. The remaining variation can be due to low editing efficiency, sampling noise or real signal of variable fitness effect of different deletions (Smits et al. 2019).

**Figure 5.**
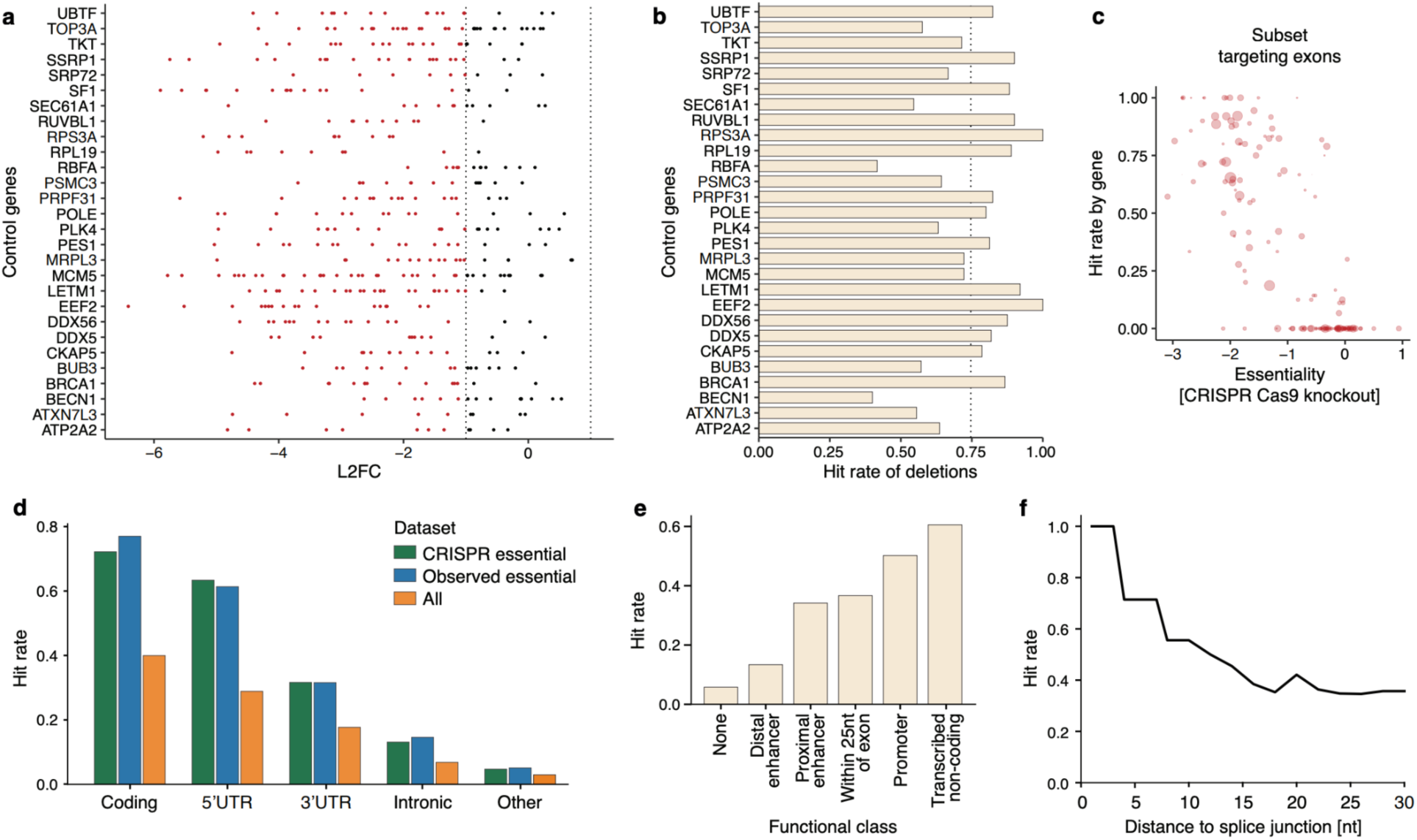
a. Log2 fold change (x-axis) for epegRNAs (markers) targeting different essential genes (external CRISPR screen LFC < -1; y-axis). Dashed line: LFC = -1. **b.** Hit rate of deletions (fraction of epegRNAs with LFC < -1, x-axis) for different essential genes (y-axis). Dashed line: 0.75. **c.** Hit rate (y-axis) of deletions targeting genes (markers) contrasted against CRISPR screen essentiality of the gene (x-axis). Marker size: number of deletions. **d.** Hit rate (fraction of deleted elements with LFC < -1, y-axis) for different gene areas (x-axis) considering CRISPR essential genes (external LFC < -1, green), observed essential genes (median coding deletion LFC < -1, blue) and all genes (orange). **e.** As (d), but for different functional classes (x-axis). **f.** As (d), but for increasing distance of deleted intronic DNA to a splice junction (x-axis).

We next explored the 462 deletions outside coding exons with strong signal (LFC < -1). First ,we asked which non-coding regions in essential genes harbor the most hits. In 5’UTR exons, 63% (128/202) of deletions dropped out, followed by 31% (93/294) in the 3’UTR, 13% (57/434) in introns, and 5% (53/1,122) in other regions, in contrast to 72% of coding region deletions (Figure 5d). This ranking of signal was consistent when focusing on genes essential in an external CRISPR screen, genes with strong average deletion fitness effect in our screen (median coding deletion LFC < -1), or all deletions in the screen. Considering functional annotations, transcribed regions (at least 30% of maximum RNA coverage of the same gene, mostly UTRs) depleted 61% (158/261) of the time when deleted, with promoter deletions also depleting most of the time (135/269, 50%), deletions of proximal enhancers depleting in 34% of the cases (181/530), and distal enhancers carrying less signal (13%; 83/621, Figure 5e). Intronic deletions had a larger impact when closer to splice junctions (Figure 5f), with all four deletions within 3nt of a junction and 56% of the nine deletions within 10nt depleting. The distance effect attenuated after ∼25nt, with 35% (11/30) of deletions showing signal. Finally, considering various computational scores of sequence importance, CADD-SV score (Kleinert and Kircher 2022) that integrates many of the annotations above predicted deletion fitness the best, with 35% (252/718) of high-scoring deletions observed depleted.

Finally, we validated the effect of individual deletions observed in the screen. First, we confirmed that deletions that did not affect cell growth are indeed viable by generating individual pegRNAs, transfecting them in an arrayed setting, genotyping the cells by PCR and Sanger sequencing. Amplified sequences from all of the 21 different pegRNA pairs were of the expected size, and encoded the deletion allele (Table S5). These viable edits include deletions of CTCF sites (Figure 6a,b), regions close to other essential deletions (Figure 6c), and conserved regions (Figure 6a-d). To validate essential deletions, we cloned 10 pairs of pegRNAs into lentiviral backbones that also encode mScarlet, and competed cells carrying them against ones expressing non-targeting control pegRNAs and BFP. Out of the 10 tested deletions (Figure 6e), seven depleted in competition compared to the controls after 11 days in culture (Figure 6f). These seven validated deletions had the strongest dropout in the screen of the tested ones (all LFC ≤ -4) , and include coding, intronic, and promoter regions. The three unvalidated ones had weaker effects (LFCs - 2.2, -1.8, -2.7) and were intergenic. Together, these results confirm both non-essential deletions that support cell viability and essential deletions that impair cell fitness, validating the utility of the paired prime editing platform for pooled functional genomic interrogation at scale.

**Figure 6.**
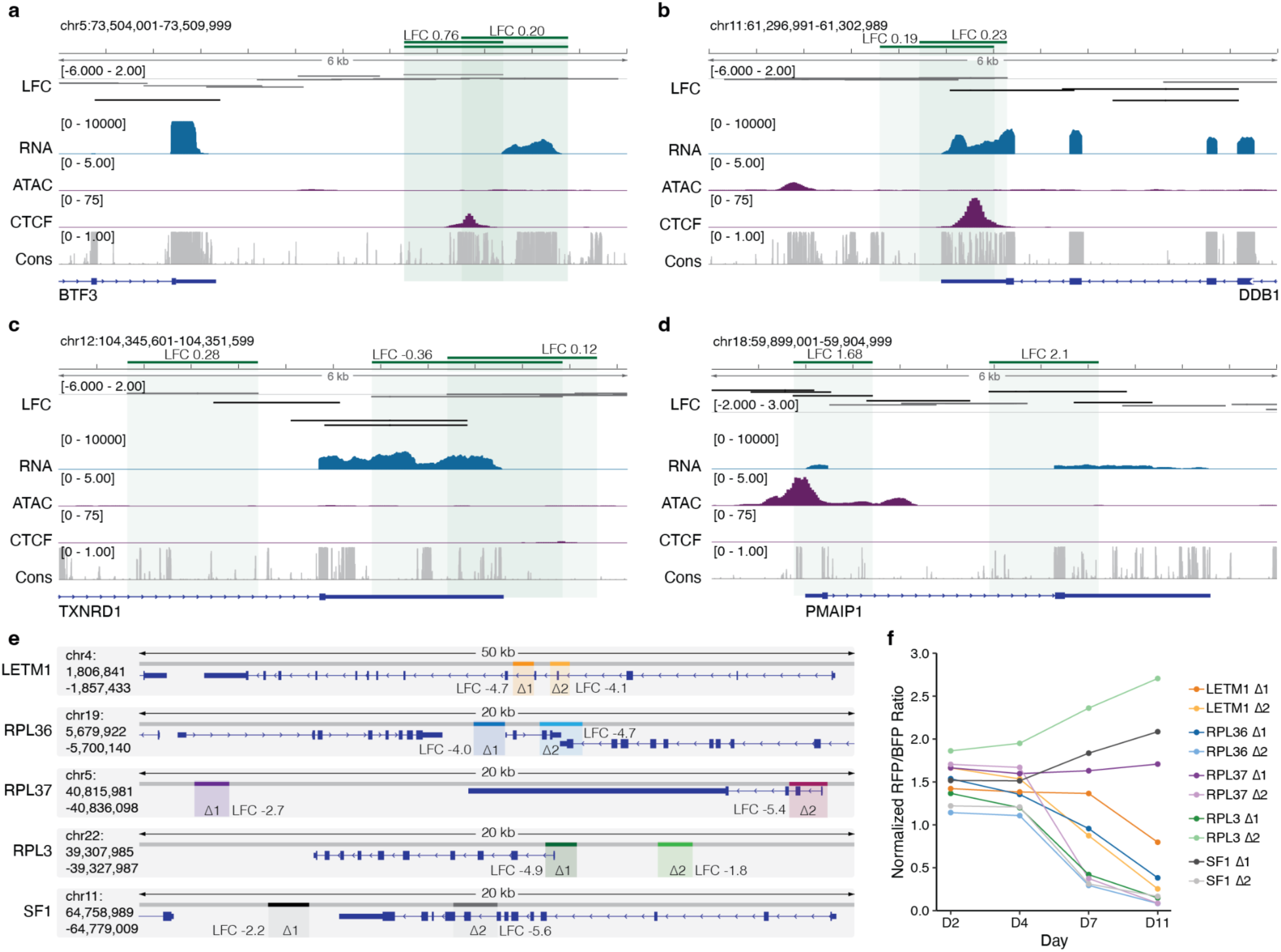
a. Genome browser visualization of validated non-essential deletions in proximity to BTF3. Green bars and shared areas: tested deletions; bar markers: signal in the screen; LFC track: log_2_-fold change in the screen, with lower values stronger effect. **b.** As (a), but for DDB1. **c.** As (a) but for TXNRD1. **d.** As (a) but for PMAIP1. **e.** Genome browser visualizations of evaluated essential deletions across five genes. **f.** Dropout of cells over time (x-axis) with targeting pegRNAs (RFP positive) standardized to non-targeting pegRNAs (BFP positive; ratio on y-axis), validating essentiality of targeted regions.

## Discussion

We developed and applied large-scale paired prime editing to generate synthetic deletions across the human genome, uncovering determinants of editing efficiency and discovering essential elements within coding and non-coding DNA. By screening thousands of epegRNA pairs, we identified deletion length, chromatin contact frequency, and pegRNA-specific features to be key in determining deletion efficiency. We used this information to demonstrate the capacity to generate clinically important deletions of up to 342 kb with high precision and at therapeutically meaningful rates. In a genome-scale pooled screen, we applied these insights to tile ∼10 kb regions around 149 essential genes in HAP1 cells, identifying hundreds of non-coding and intronic regions with fitness effects. Our results underscore the maturing potential of prime editing for functional interrogation on the scale of the entire genome.

Our findings highlight the advantages of paired prime editing in achieving scarless and precise deletions without introducing double-strand breaks. Whereas paired Cas9-based methods typically yield ∼10-50% accuracy at deletion junctions (Guo et al. 2018; Ousterout et al. 2015) and are associated with significant off-target effects and chromosomal instability (Kosicki, Tomberg, and Bradley 2018), the paired prime editing deletions were generated with around 95% purity at the junctions and at high efficiency thanks to the PE7 system (Yan et al. 2024). This aligns with prior reports in a limited number of loci (Anzalone et al. 2019), but extends the learnings to high-throughput pooled applications. Unlike recombinase-based methods, which require pre-installed recognition sites, our method works on native sequences, making it applicable for genome-scale functional studies and potential one-shot therapeutic interventions. Given our conservative estimate of ∼70% recall of hits, and assuming a tiling design of 3-fold coverage of any nucleotide, as well as independent editing efficiencies of individual pairs, the false negative rate of pooled deletion screens is expected to be less than 3%.

The implications of the improved ability to precisely and scalably delete kilobases of DNA span several domains. First, we provide a broadly applicable platform for assessing the essentiality of non-coding DNA in a pooled screen approach, showing that most intergenic and intronic sequences near essential genes are dispensable for growth in HAP1 cells. However, a notable subset enriched in proximity to transcription start sites and splice junctions does impact cell fitness. Together with improved DNA synthesis technologies that now enable library designs with both pegRNAs on a single construct, the pooled deletion approach opens avenues for dissecting enhancer function, gene regulation, and the roles of long non-coding RNAs. Second, the ability to engineer large deletions reliably highlights the potential of paired prime editing for therapeutic genome editing. For example, we could achieve therapeutically meaningful deletions in genes such as retinitis pigmentosa-linked *RHO* and Duchenne muscular dystrophy causing *DMD*, offering a path to replace or bypass mutation-rich exons through deletions as large as 340 kb. Third, the predictive model we developed explained up to 60% of the variance in editing efficiency, adding valuable input for pegRNA design that can guide users toward combinations of high-efficiency sites. Deletion length and contact frequency emerged as the strongest predictors, with predicted individual pegRNA efficacies also playing a role.

Some of our experimental choices limit the generalizability of our observations. The initial reporter-based deletion screen was restricted to the mScarlet reporter locus on chromosome 13, which resulted in confounding between deletion size and chromatin context, potentially biasing the results. Furthermore, only the forward-strand-targeting epegRNA was varied, precluding exploration of guide-pair interactions. In the pooled genome-scale screen, we found three genes (*EIF4B*, *EIF4G1*, *POLG2*) that strongly and reproducibly deviated from the observed CRISPR data. Some genes, e.g. *TRRAP* with 72 coding exons, were deleterious in the CRISPR screen, but showed less signal upon deletion of some individual exons, potentially due to natural tolerance to splicing errors. Indeed, frame preserving deletions in this case were substantially more tolerated compared to frame disrupting ones (average LFC 0.19 vs -0.53). These findings suggest biological, rather than technical reasons for some unexpected signal (Smits et al. 2019), and imply that our false negative estimates may be conservative.

Overall, we demonstrated the utility of paired prime editing for scalable, high-resolution genome interrogation. We provided the first genome-scale deletion screens using paired pegRNAs, uncovering editing efficiency determinants and functional non-coding sequences. In the future, integrating single-cell transcriptomics, extending the approach to disease-relevant cell types, and improving prediction models will further broaden the scope of such screens. Ultimately, paired prime editing promises to unlock functional secrets from all the gigabases of our genome.

## Author contributions

Designed experiments: JW, GG, EMP, SK, LP. Performed experiments: JW, GG, EMP, YB. Analyzed data: JW, JK. Provided technical guidance: FGL, SI, TE. Wrote paper: JW, LP with input from all authors.

## Acknowledgements

All authors were supported by Wellcome (220540/Z/20/A). We thank members of the Parts lab for input on the text, and Britt Adamson for sharing PE7 plasmids.

## Data availability

The measurements from experiments are shared as supplementary tables.

## Code availability

Code for paper results is at https://github.com/julianeweller/PrimeDeletions

## Methods

### Mammalian cell culture

Cells were cultured at 37 °C and 5% CO2, with HAP1 in IMDM (Invitrogen) and HEK293T cells in DMEM (Invitrogen), both supplemented with 10% FBS (Invitrogen), 2 mM glutamine (Invitrogen), 100 U/ml penicillin and 100 mg/ml streptomycin (Invitrogen). For splitting, cells were washed with PBS (Sigma Aldrich) and dissociated with 0.5x TrypLE Express (Gibco). After 5 min incubation at 37 °C, TrypLE was inactivated with culture media, and cells were spun down for 3 min at 350 to remove the supernatant and plate them into a new culture vessel. For cryopreservation, cells were resuspended in 1 ml of culture media supplemented with 10% DMSO (Sigma Aldrich), frozen at -80 °C using MrFrosties (Nalgene), and then transferred to liquid nitrogen.

### Plasmid cloning

pegRNAs used at different steps are provided in Table S6, and references for relevant plasmid sequences are in Table S7. The donor plasmid Hygro-BFP-attP was ordered from Twist Biosciences for integration into the attB site with GT central dinucleotides using Bxb1 recombinase. Central dinucleotides were mutated to AC using Q5 Site-Directed Mutagenesis Kit (NEB) following the manufacturer’s instructions. The plasmids wtBxb1 (Addgene #222337), evoBxb1 (Addgene #222338), and eeBxb1 (Addgene #222339) were a gift from David Liu. The epegRNAs were cloned by assembling the spacer, scaffold, and 3’ extension into the pU6-epegRNA-GG-acceptor backbone as described in Anzalone et al. (Anzalone et al. 2019). Forward and reverse oligonucleotides were ordered from IDT and hybridized, creating overhangs matching the other components or the integration site in the backbone digested with BsaI. The components were assembled in a single reaction using Golden Gate assembly and transformed into XL10 gold ultracompetent bacteria (Agilent).

### Cell line generation

To generate the HAP1-PE2-mScarlet reporter cell line with mScarlet in the CLYBL locus, the HAP1-PE2 cell line was transfected with 1 µg pCLYBL_CAGG_mScarlet, 1 µg of pCAG-Cas9-U6-CLYBLgRNA, and 210 ng of a GFP-puromycin plasmid using Xfect (Takara Bio) following supplier’s instructions. For the HEK203T-PE2-mScarlet reporter cell line with mScarlet-degron in the CLYBL locus, the HEK293T-PB-PE2 cell line was transfected with 1.2 µg pCLYBL_CAGG_mScarlet-degron, 1.2 µg pCAG-Cas9-U6-CLYBLgRNA, and 300 ng of a GFP-puromycin plasmid using Lipofectamine LTX (Invitrogen) following supplier’s instructions. Both cell lines were selected with puromycin for 3 days. 2 weeks post-transfection, cells were sorted into single clones based on GFP (prime editor) and mScarlet (reporter) expression.

For generating the HAP1-PE7 cell line, the plasmid pLenti_PE7-GFP was generously provided by Britt Adamson. Lentivirus was produced by transfecting 0.85 million Lenti-X HEK293 cells (Takara) seeded in 6-well plates with 10 µg pLenti_PE7-GFP, 3.8 µg pMD2.G (Addgene 12259), and 7 µg psPAX2 (Addgene 12260) in 250 µl OPTI-MEM and 6 µl Trans-IT-LT1 (Mirus Bio). After a 15-minute incubation, the transfection mix was applied to the cells. ViralBoost Reagent (Alstem) was added at 2× after 6 h. The lentivirus-containing medium was collected after two days, filtered through 0.45 µm cellulose acetate filters (VWR), and titrated with Lenti-X GoStix Plus (Takara). The HAP1 ΔMLH1 cell line was purchased from Horizon Discovery (HZGHC000343c022). It was transduced with the PE7-GFP lentivirus using a final concentration of 8 µg/ml polybrene and spun for 1h at 1,000×g at room temperature. Two days after transduction, GFP-positive cells were bulk-sorted, and a week later, they were single-cell cloned. Clones were assessed for GFP levels using flow cytometry and clones with high GFP levels were expanded and characterized.

### Mutation rate analysis for editing outcomes

Paired reads were trimmed to the target site using 20bp adaptors adjacent to the target sites with cutadapt v4 (Martin 2011) using a minimum length of 20 and a quality cutoff of 15. They were then paired with PEAR v0.9.11 (J. Zhang et al. 2014). A custom R script was used to generate count tables of editing outcomes. In short, reads were grouped by sequence and counted. Then, the sequence between the target sites was extracted to identify the inserted sequence and compare it to the intended attB site.

### Library design

For the anchored library, first, all targets with NGG PAM were extracted from the 2 Mb region surrounding the reporter locus on Chromosome 13. All spacer candidates were scored using FlashFry (McKenna and Shendure 2018) and DeepSpCas9 (Kim et al. 2019). Spacers with non-ATGC characters in the target site, polyT, more than three matches to the reference genome, a CFD score (Doench et al. 2016) below 0.01, or a DeepCas9 (Kim et al. 2019) score below 10 were removed. The remaining spacer candidates were categorized by length bins with the boundaries 1kb, 5kb, 10kb, 50kb, 100kb, 500kb, 1 Mb, and longer. At least 250 spacers or all available spacers for each bin were selected based on the highest DeepSpCas9×CFD score. All spacers were combined with an 11nt PBS, but a small subset was also screened with 9nt and 13nt. All spacers were combined with a 34nt homology arm carrying a subset of the attB attachment site. Additionally, a small subset with homology arm modifications were also screened. Central AC dinucleotides were chosen for the attB site. The main library uses the improved epegRNA scaffold (GTTTAAGAGCTATGCTGGAAACAGCATAGCAAGTTTAAATAAGGCTAGTCCGTTATCAACTTGAAAAAGTGGCACCGAGTCGGTGC). Additionally, epegRNAs with the PASTE scaffold (GTTTGAGAGCTATGCTGGAAACAGCATAGCAAGTTCAAATAAGGCTAGTCCGTTATCAACTTGAAAAAGTGGCACCGAGTCGGTGC) (Yarnall et al. 2023) were included. The sequence was assembled and endowed with cloning sites for BsaI. The scaffold was integrated in a secondary cloning step using BsmBI sites.

For the paired deletion library, 149 genes were selected based on gene expression and essentiality for HAP1 cell lines. All spacers in the selected genes were extracted using the WGE CRISPR design tool (Hodgkins et al. 2015) and filtered for 20-80% GC content. The spacers were scored using DeepCas9 (Xue et al. 2019), and only those with scores exceeding the mean minus one standard deviation were retained. Off-targets were scored with WGE and filtered for no off-targets with 0 or 1 distance, less than 4 off-targets with a distance of 2. For the remaining spacers, pegRNAs were assembled and scored with PRIDICT2 and ePRIDICT (Mathis et al. 2024). Since PRIDICT2 can only score single pegRNAs with an RTT sequence homologous to the target site, the scores correspond to the predicted insertion efficiency of the homology arm into the target site as an approximation. Guides were retained if their PRIDICT2 scores exceeded the mean minus one standard deviation for both the HEK and K562 model scores, while a cutoff of 30 was applied for ePRIDICT scores.

To generate deletions, a forward-strand targeting pegRNA was paired with a reverse-strand targeting pegRNA further downstream. This was achieved by iterating through each gene, selecting all available spacers in the gene and its surrounding region (+/- 11 kb) and choosing spacers based on deletion length and predicted efficiency. The final deletions ranged from 298 bp to 4,243 bp (median = 1,098 bp). Finally, 240 non-targeting controls randomly sampled from the Brunello library subset were selected and paired with each other, leading to 120 paired guides (Doench et al. 2016). The first nucleotide in each spacer was replaced with a G for improved expression of the guide. PBS length was set to 13 nt and homology arms to 34 nt, carrying the attB overhang sequences (TCCTGACGACGGAGGTCGCCGTCGTCGACAAGCC, TGTCGACGACGGCGACCTCCGTCGTCAGGATCAT). The oligos were assembled containing the following components sequentially: primer site (CCTCTGGTCGGATGA), BsaI site (GGTCTCCCACC), pegRNA1 (guide + improved scaffold + attB overhang + PBS), cloning site (CGCGGTGAGACGGTTTAAACCGTCTCCCTTG), pegRNA2 (guide + cr772 scaffold + attB overhang + PBS), BsaI site (CTACGTGAGACC), and another primer site (TGGGTCCTCAGATCT). Five oligonucleotides with intrinsic BsaI sites were removed. The final library contained 11,084 epegRNA pairs.

### Library cloning

The designed anchored library was synthesized as an oligonucleotide pool (Twist Biosciences), amplified for 16 cycles with KAPA HiFi HotStart ReadyMix (Roche) and purified with the Monarch PCR & DNA Cleanup Kit (NEB). To clone the library into the lentiviral acceptor backbone pLenti_epeg_acceptor, 2.1 ng of the pool and 67.75 ng of the backbone (1:3 molar ratio of insert and vector) were digested with BsmBI-v2 (NEB) and purified using the Monarch PCR & DNA Cleanup Kit. The components were combined with Golden Gate assembly, including BsmBI-v2 in the reaction with T4 DNA ligase (NEB), cycling between 5 min at 16°C and 42°C for 30 iterations and a final heat inactivation step at 60°C. The ligation products were purified with the Monarch PCR & DNA Cleanup Kit, and ten electroporations were performed into MegaX DH10B T1R Electrocomp Cells (Thermo Fisher). Library coverage at this step was > 100 based on colony counts on plates of the library in comparison with religation controls. The bacteria were grown in a liquid culture overnight, and plasmid DNA was extracted using the Plasmid Plus Midi Kit (Qiagen). The plasmid sequence was first verified with Sanger sequencing and then sequenced on the Illumina MiSeq 250PE platform as described below.

The paired deletion library (ordered from Twist Biosciences) was amplified for 6 cycles with the KAPA polymerase (Roche) using 0.5ng as input for each of the 8 PCR reactions. It was then digested with BsaI (NEB) for 1 h at 37 °C, followed by a heat inactivation step at 80 °C for 20 min. The backbone plasmid pLenti-puro-mCherry-epeg was digested with BsmBI-v2 for 2 h at 55 °C and heat-inactivated. Both products were purified using the Monarch PCR & DNA Cleanup Kit (NEB). T4 ligase (NEB) was used for ligation at a 3:1 molar ratio of insert to backbone at room temperature for 1 h, followed by heat inactivation. The product was electroporated into XL-1 Blue Competent Cells (Agilent) and extracted with the QIAGEN Plasmid Plus Midi Kit. In a secondary cloning step, gene fragment GF1 containing the tevopreQ motif and the terminator for the first epegRNA, and a mU6 promoter for the second epegRNA were inserted into the cloning site of the library. The library was digested with BsmBI-v2 for 1 h before setting up the Golden Gate reaction for 99 cycles with a 2:1 ratio of insert:backbone, the T4 ligase, and BsmBI-v2 (NEB). The product was purified with the Monarch DNA & PCR clean-up kit (NEB) and electroporated into XL-1 Blue Competent Cells (Agilent). The final library was extracted with the QIAGEN Plasmid Plus Midi Kit.

### Lentivirus production

To produce lentivirus in HEK293FT cells, 32.4 µg of the lentiviral library plasmid pool was mixed with 32.4 μg of psPax2 (Addgene 12260) and 7.2 μg of pMD2.G (Addgene 12259) in Opti-MEM (Gibco) together with 72 µl of PLUS reagent (Invitrogen). After 5 min of incubation at room temperature, 216 µl of LTX (Invitrogen) was added, and the mix was further incubated for 30 min. Then, 3 ml were dropped onto 10-cm dishes with 80% confluent cells in DMEM medium. The supernatant containing the virus was collected after 48h and 72h. The lentiviral titer was tested with Lenti-X GoStix Plus (Takara) showing a strong positive signal. To determine the volume of virus required to reach a multiplicity of infection of 0.3, the virus was transduced into 12-well and 96-well plates, and cell viability was quantified for cells under selection with 2 µg/ml puromycin and without selection using the CellTiter 96 Aqueous One Solution Cell Proliferation Assay (Promega) following the instructions from the manufacturer.

### Anchored deletion screen

The HEK293T and HAP1 reporter cell lines expressing mScarlet in CLYBL and inducible prime editor PE2 were transduced with the virus pool, aiming for a multiplicity of infection of 0.3 and an epegRNA coverage of >1,000x. For each replicate, 13.4M cells were spin-infected in 4 wells of a 6-well plate for 30 min at 1,000g. Cells were resuspended and plated at 2×10^4^ cells per cm^2^. The next day, puromycin was added at 2 µg/ml to select for library integration for a minimum of 3 days. The prime editor was induced with 1 µg/ml doxycycline 1h before transfection of reverse-targeting anchor epegRNA. For HAP1 cells, the reverse-targeting anchor epegRNA was nucleofected into the library cells with the Neon Transfection System (Invitrogen) following the manufacturer’s instructions, assuming a 25% nucleofection rate to aim for >1,000x coverage. For HEK293T, the anchor epegRNA was delivered with lipofection using Lipofectamine LTX (Invitrogen), assuming a 50% transfection rate to aim for >1,000x coverage. The media was replaced the next day, and 1 µg/ml doxycycline and 10 µg/ml blasticidin were added to express the prime editor and select for transfected cells. The screens were maintained at 2,000x coverage for two weeks, with 7 days of induced prime editor. On the day of harvest, screens were sorted into mScarlet-positive and negative populations using fluorescence-activated cell sorting. To ensure coverage of >2,000x, 8.8M - 11M cells per screen were sorted. On average, around 11% of the cells were negative for mScarlet. Each screen was performed in at least two replicates. For each sorted cell population and screen, gDNA was extracted from 5M cells using the DNeasy Blood & Tissue Kit (QIAGEN) with the addition of RNase (Thermo Fisher). The sequencing libraries were prepared as described below for sequencing on Illumina MiSeq 250 bp paired-end with 36 dark cycles on read 1, and Illumina NovaSeq SP 150bp paired-end platforms.

### Paired deletion screen

The HAP1-PE7 cells were spin-infected for 30 min at 1,000 xg with the paired epegRNA library, aiming at a multiplicity of infection of 0.3 and a guide coverage of >2,000x. The screen was performed in two replicates that were independently infected. Cells were cultured in T875 flasks (Corning), and cells were selected for library integration after 18h using 2 µg/ ml puromycin for 3 days. Cells were split and pelleted every 3-4 days, from day 4 to day 28. Expression of GFP for the prime editor and mCherry for the library were monitored weekly. Additionally, on day 21, attP donor Hygro-BFP-attP (AC dinucleotides) and evoBxb1 were delivered to a subset of cells from screen replicate 2, corresponding to 4,000x library coverage, using the Neon NxT Transfection System (Invitrogen) following the manufacturer’s instructions. They were cultured for another 14 days under 0.15 mg/ml hygromycin selection. On day 32, a subset of the screen with Bxb1 integration was sorted for BFP-positive cells with the Bigfoot Spectral Cell Sorter (Thermo Fisher) and also harvested on day 35. The genomic DNA was extracted from the cell pellets for each replicate and time point, as described by Allen et al. (Allen et al. 2018).

### Library preparation for next-generation sequencing

The epegRNA constructs of the anchored library were amplified with a mix of forward primers (ACACTCTTTCCCTACACGACGCTCTTCCGATCTGATGGCTTTATATATCTTGTGGAAAGGA CGAAACACC and ACACTCTTTCCCTACACGACGCTCTTCCGATCTCTAGAATGGCTTTATATATCTTGTGGAAAGGACGAAACACC) and the reverse primer (GTGACTGGAGTTCAGACGTCTGCTCTTCCGATCTTGACCGCTGAAGTACAAGTGGT) using Q5 HotStart High-Fidelity 2X Master Mix (NEB). The amplicons were column-purified using the QIAquick PCR purification kit (QIAGEN) and prepared for Illumina sequencing with KAPA HiFi HotStart ReadyMix (Roche) and indexing primers for 14 cycles and with an annealing temperature of 72°C. The PCR products were purified with AMPure XP beads (Agencourt) in a 0.7:1 ratio (beads to PCR reaction volume). The samples were quantified on the Bioanalyzer (Agilent) with the DNA High Sensitivity kit and pooled for sequencing on Illumina platforms.

For the paired deletion screen, to maintain >2,000x coverage, gDNA corresponding to 25 million cells per screen was used to amplify the epegRNA locus by PCR using repliQa HiFi ToughMix (QuantaBio). To increase the diversity at each sequencing position, 4 staggered forward primers were used. Each PCR reaction was run in 50 µl reaction volumes with no more than 4 µg of input DNA and run for 20 cycles. The products were purified with the QIAquick PCR Purification Kit (Qiagen). In a second PCR with 12 cycles using KAPA HiFi HotStart ReadyMix (Roche), the sequencing adaptors and barcodes are added to 1ng input DNA. Each sample was indexed with a distinct pool of up to indexes. The amplicons were purified with AMPure beans using a 0.7:1 ratio of beads to PCR reaction volume. The amplicons were pooled and sequenced on Element AVITI using the 2×300 CB Freestyle High Output Kit (600 cycles, 300 paired-end).

### Analysis of the paired deletion screen

To pair the forward and reverse reads, the reverse complement of the reverse read was generated with BBMap v39 (Bushnell 2014). The reads were then stitched together using a custom Python script, filtered to contain both spacers and primer binding sites, and counted for pegRNA pairs. One base pair mutations, assumed due to errors in PCR or sequencing, were corrected if the components matched a unique library sequence with a maximum one base pair distance. For each time point and epegRNA pair, their frequencies were calculated by dividing their count by the total read counts for this sample. The log2 fold change (LFC) for day 14 and day 28 compared to day 4 was then calculated by subtracting the log2 value of the day 4 frequency from the log2 value from the day 14 or day 28 frequency. For further analysis, the mean of the LFC of day 11, day 14, day 21, and day 28 was used. For each epegRNA pair, the frequencies at which they had BFP integrated (“BFP signal”) were calculated by dividing their sequence count in the sorted BFP positive cell population by the total read counts for this sample and then averaged across sequencing and biological replicates.

Features described during library design were added to the read counts. Further, gene annotations were downloaded from GENCODE (release 47, GRCh38.p14) (Frankish et al. 2023). The transcription start site positions were downloaded from the UCSC Genome Browser (Perez et al. 2024). Transcripts were filtered for genes targeted in the library. The first transcript start and the last transcript end were used to define the distance to the transcription start and end positions. The features were assigned to epegRNA pairs using the bedtools2 intersect function with options -wa -wb (Quinlan and Hall 2010). The read counts were visualized in a custom R script.

Protein-coding gene, transcript, exon, and untranslated region (UTR) annotations for the human genome were obtained from the Ensembl database using the biomaRt R package (dataset: hsapiens_gene_ensembl). Genomic coordinates, Ensembl gene identifiers, HGNC symbols, and strand information were retrieved for all protein-coding genes. Transcript-level data were then downloaded, and only those annotated as principal by the APPRIS database were retained. Exon coordinates corresponding to these APPRIS principal transcripts were extracted. RNA-seq coverage data from HAP1 cells were imported using the rtracklayer package, and exon-level CPM values were calculated by overlapping RNA-seq signal with exon coordinates. Exons with CPM ≥10 were defined as transcriptionally active. These expressed exonic intervals were intersected with protein-coding gene coordinates to retain only those belonging to coding genes. Exons were subsequently ranked according to their genomic position relative to the transcription start site, accounting for strand orientation. Finally, 5′ and 3′ untranslated regions (UTRs) were annotated by identifying overlaps with the ranked exon set using the findOverlaps function from the GenomicRanges package. Each exon was classified as part of a 5′-UTR, 3′-UTR, or coding region, resulting in a final dataset comprising 173,680 exons across 14,580 genes.

### Flow cytometry

For analysis, the CytoFLEX Flow Cytometer (Beckman) was used with the CytExpert software and FlowJo Version 10. Events were gated for cells based on forward scatter (FSC-H) and side scatter (SSC-H), single cells were isolated from forward scatter (FSC-A) and its width (FSC-W). GFP was analyzed in the channel 525/40 (488nm) (FL5-A), mScarlet in 610/20 (561nm) (FL8-A), and BFP in 450/45 (405nm) (FL1-A). Sorting was performed with 100µm nozzles on either Sony MA900, Sony SH800S, or by the flow cytometry core facility on MoFlo XDP (Beckman Coulter), Bigfoot Spectral Cell Sorter (Thermo Fisher), or BD Influx Cell Sorter (BD Bioscience).

### Targeted long-read nanopore sequencing

Targeted long-read sequencing was performed on Nanopore MinION with R9.4.1 flow cell following the manufacturer’s protocol for Ligation sequencing gDNA - Cas9 enrichment (SQK-LSK109). The gDNA was extracted with the HMW DNA Extraction kit (Monarch), and 5 µg was used for sample preparation. The gDNA was dephosphorylated for 20 minutes and then cleaved with custom-designed guides (TTTAAGATCCTGGCAGCAAGAGG , AGAGCTCTATTGGAAATCAGTGG, TGCGATCTGCTGTCTGCAAACGG, GCACTGGTTAGATTTGGCAGAGG). The guides were designed with CHOPCHOP (Labun et al. 2019) and ordered from IdT as SpCas9 Alt-R crRNAs together with SpCas9 Alt-R tracrRNA and Aly-T Sp HiFi Cas9 nuclease V3 (IDT) to form Cas9 RNPs. After dA-tailing and adapter ligation, the sample was purified with AMPure XP beads (Beckman Coulter) and eluted overnight. The library was loaded onto the flow cell and sequenced for 24 - 48 h. The pod5 results files were aligned to the reference genome with Dorado 0.8 using the dna_r9.4.1_e8_sup@v3.6 model and visualized in the IGV Genome Browser (Robinson et al. 2011).

### RNA sequencing

RNA was extracted from 5 million cells per sample using the RNeasy plus mini kit (Qiagen) and prepared for sequencing using the NEBNext Ultra II Directional RNA Library Prep Kit for Illumina (NEB). The samples were pooled and run on one lane with NovaSeq S4 using 150nt paired-end sequencing. Transcripts were quantified using the quant pipeline from Salmon 1.10 against the default reference index. Downstream analysis was performed using custom R scripts to map transcripts to genes using tximport version 3.20 and visualize counts.

### Micro Capture-C

Micro Capture-C (MCC) measures three-dimensional contacts of a specified region with the genome at base pair resolution (Hua et al. 2021). To generate interaction scores between the region targeted by the reverse-anchor epegRNA in the deletion screen and the library epegRNAs, MCC was performed on the reporter cell lines with two biotinylated IDT xGen Custom Lockdown Probes (IDT) targeting the end of the mScarlet reporter (TCCGAGGGCCGCCACTCCACCGGCGGCATGGACGAGCTGTACAAGTCCGGACTCAGATCTCGAGCTCAAGCTTCGAATTCTGCAGTCGACGGTACCGCGGGCCCGGGATCCACCGGATCT, GACATCACCTCCCACAACGAGGACTACACCGTGGTGGAACAGTACGAACGCTCCGAGGGC CGCCACTCCACCGGCGGCATGGACGAGCTGTACAAGTCCGGACTCAGATCTCGAGCTCAA) following the steps detailed in the published protocol from Hamley et al. (Hamley et al. 2023).

In summary, cell pellets of the reporter cell lines were fixed in formaldehyde (Sigma-Aldrich) at 2% final concentration, which was quenched with 15% final volume 1 M glycine (Sigma-Aldrich). They were then pelleted and frozen at -80 °C. In the first batch, the HEK293T sample and one HAP1 sample were digested with 10 Kunitz units of MNase (NEB). In the second batch, three HAP1 reporter cell line samples were digested, with 7.5, 10, and 15 Kunitz Units of MNase each. Then, ligation was performed using the T4 DNA Ligase (NEB) and the Klenow DNA Pol I Large Fragment (NEB). To prepare the sequencing library, the first batch was sonicated on an ME220 sonicator (Covaris) for 80-90s and the second for 180 - 335s. Next, end prep, adaptor ligation, and indices addition were performed with the NEBNext Ultra II Library Kit (NEB). Finally, two rounds of capture enrichment with the HyperCapture Target Enrichment Kit (Roche) were conducted with 44h - 70h and 24h incubation periods. The first batch was sequenced on Illumina MiSeq 150PE, probe 1 of the second batch was sequenced on Illumina MiSeq 150PE, and probe 2 on Illumina NovaSeq 150PE.

The data was analyzed based on the steps outlined in Hamley et al. (Hamley et al. 2023). First, adaptors were removed using Trim Galore (Version 0.6.5). Overlapping reads were merged with FLASH (Version 1.2.11) (Magoč and Salzberg 2011) with a maximum overlap of 150. The reads are then mapped to the regions surrounding the probes with Blat using a minimum score of 20, a minimum identity value of 5, a maximum intron value of 10,000, and a tile size of 11. The reads are split using the MCC_splitter.pl script kindly provided by Oxford University Innovation Limited and aligned to the reference genome using bowtie2 (version 2.5). The resulting files are analyzed with the MCC_analyzer.pl script kindly provided by Oxford University Innovation Limited and converted to bam files with samtools (version 1.21) and bigwig files with bedtools (Version 2.21).

### Predictive modeling

The anchored deletion dataset with 11nt PBS and 34bp homology arm was split into a train and test set at a ratio of 0.7 for training. Hyperparameters were tuned for the XGBoost model (Chen and Guestrin 2016) by evaluating the average model performance based on the Root Mean Squared Error (RMSE) after five-fold cross-validation using each combination of hyperparameters (eta: 0.01, 0.05, 0.1; max_depth: 4, 6, 8; subsample: 0.6, 0.8, 1.0; colsample_bytree: 0.6, 0.8, 1.0; nrounds: 400, 500, 600). For the final model, a learning rate of 0.05, a maximum depth of 6, a subsample rate of 0.8, and a subsample ratio of columns when constructing each tree of 0.8 were used for 400 rounds, optimizing for RMSE. The selected features are listed in Table S3. Predictions were evaluated using Pearson’s correlation coefficient (R) and the variance explained (R2). The SHapley Additive exPlanations (SHAP) values and feature importance were calculated using the SHAP TreeExplainer (Lundberg and Lee 2017).

### Software and packages

IGV 2.18.4, PRIDICT2.0, ePRIDICT, DeepCas9, WGE CRISPR design tool, BBMap v39, bedtools 2.31, FLASH 1.2.11, samtools 1.21, bowtie2 2.5, CHOPCHOP, FlashFry, DeepCas9, bigWigToBedGraph v469

Python: python 3.8, pandas 2.0.3, numpy 1.24.4, matplotlib 3.7.3, seaborn 0.13.2, biopython 1.83, genet 0.14.1, regex 2024.11.6, mpmath 1.3.0, sklearn 1.3.2, xgboost 2.0.3, scipy 1.10.1, shap 0.44.1

R: R 4.4.2, tidyverse_2.0.0, ggplot2_3.5.1, tibble_3.2.1, tidyr_1.3.1, dplyr_1.1.4, stringr_1.5.1, readr_2.1.5, caret 7.0.1, xgboost 1.7.8.1, ggpointdensity 0.1.0, viridis 0.6.5, egg 0.4.5, grid 4.4.2, scales 1.3.0, reshape2 1.4.4, rtracklayer_1.64.0, ggridges_0.5.6, Gviz_1.48.0, GenomicRanges_1.56.2

