## Supplementary Material for "Generating long deletions across the genome with pooled paired prime editing screens"

### Supplementary Figures

Figure S1: Design of deletion efficiency screen via twinPE at mScarlet reporter locus.
Figure S2: Anchored deletion screen statistics.
Figure S3: Targeted long-read sequencing of mScarlet-negative cell populations.
Figure S4: Editing outcomes at the twinPE deletion junction.
Figure S5: Micro-capture C workflow, reproducibility, and correlation with screen features.
Figure S6: HAP1-PE7 cell line generation.
Figure S7: Correlation of replicates and time points in the paired epegRNA screen.
Figure S8: Correlates of log-fold changes in the paired epegRNA screen

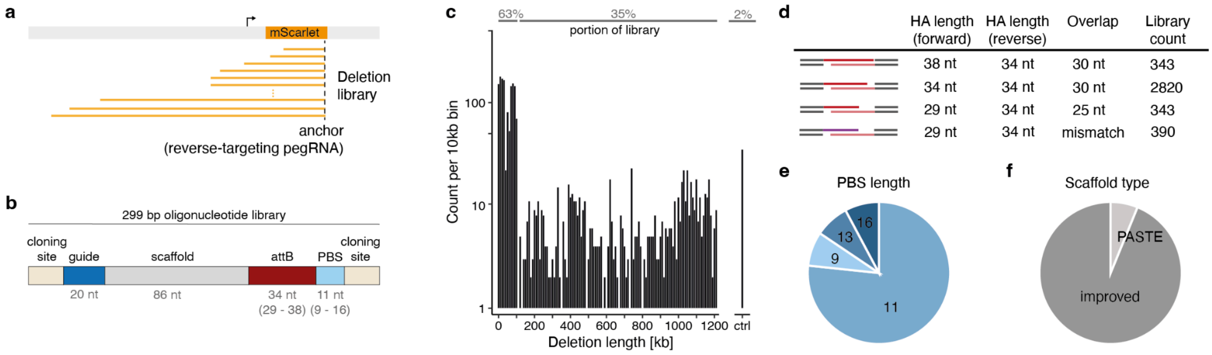


**Figure S1: Design of deletion efficiency screen via twinPE at mScarlet reporter locus.** **a.** Schematic screening approach with epegRNA anchor and various deletion lengths generated by forward-targeting epegRNA library. **b.** Schematic representation of oligonucleotide library composition with two cloning sites that carry PCR handles and BsmBI restriction enzyme sites and the epegRNA components: guide, scaffold, attB site as homology arm, and primer binding site (PBS). **c.** Distribution of deletion lengths across the library as counts per 10 kb length bins. Top: portion of the library creating different length intervals. **d.** Table summarizing used homology arms and the resulting overlap when paired with a 34-nt homology arm from anchor epegRNA. **e.** Pie chart of PBS lengths in the library, with 11 nt being the most common. **f.** Pie chart of the scaffold type, including the improved and PASTE scaffold designs.

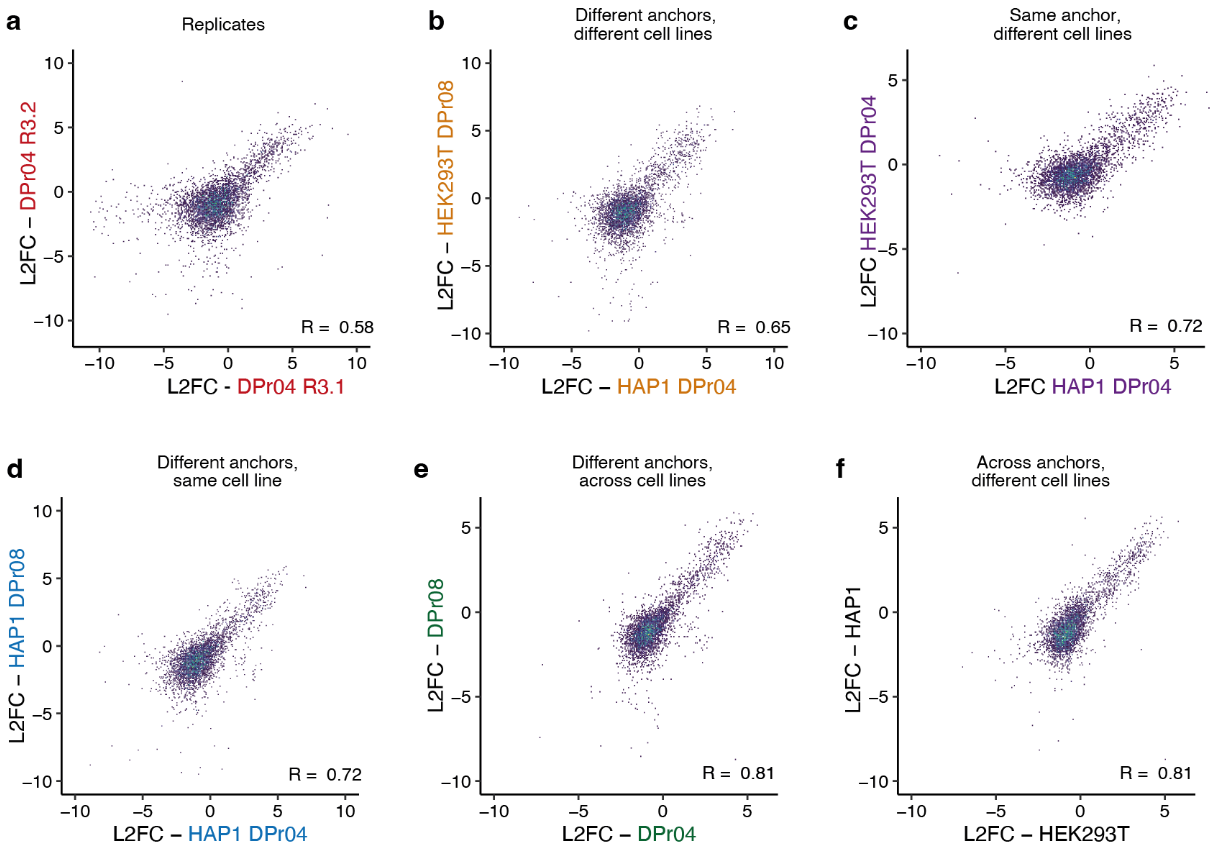


**Figure S2: Anchored deletion screen statistics. a-f.** Log2 fold changes (L2FC, x- and y-axis) of individual library members as examples for replicates (a), different anchors in different cell lines (b), same anchor in different cell lines (c), different anchors in same cell line (d), different anchors across cell lines (e), and across anchors in different cell lines (f).

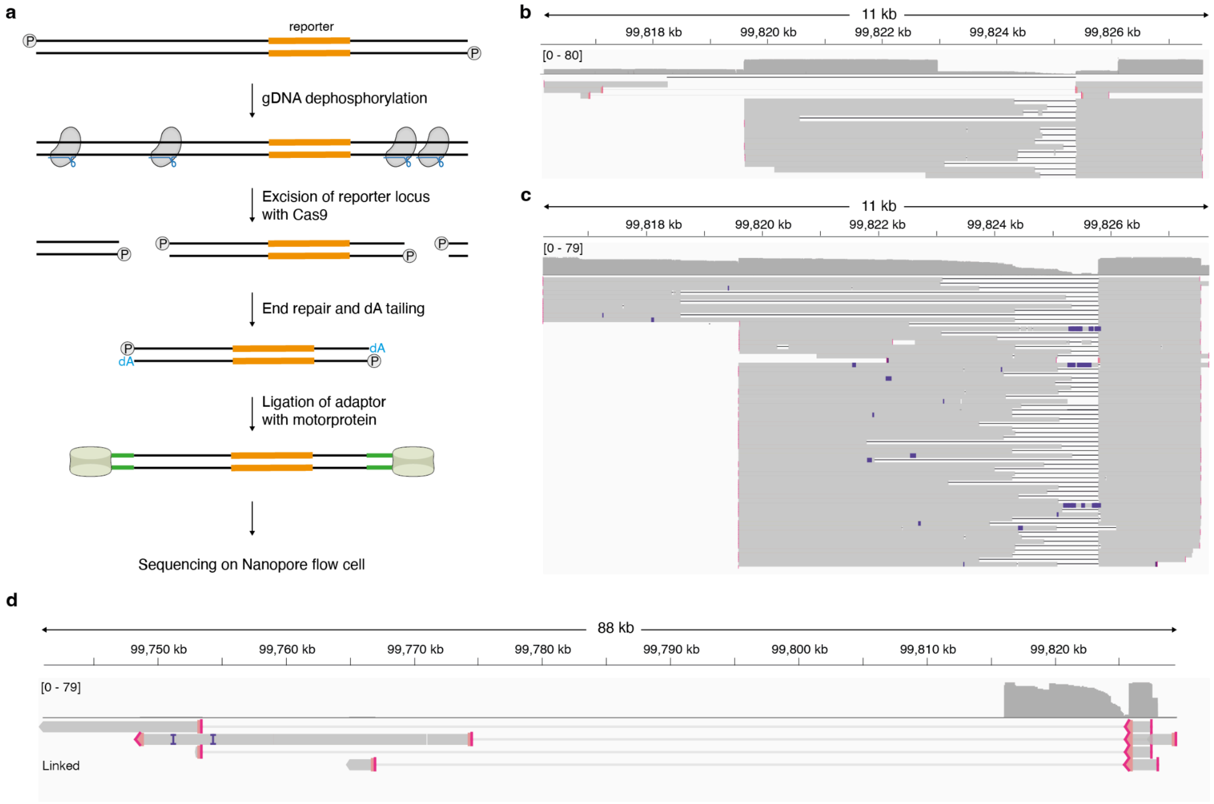


**Figure S3: Targeted long-read sequencing of mScarlet-negative cell populations. a.** Schematic representation of library preparation workflow. gDNA: genomic DNA. **b.** Read alignments from the HEK293T reporter cell line screen using the anchor epegRNA R2. Only reads aligning to reads covering the reporter are shown. **c.** Read alignments from the HAP1 reporter cell line screen using the anchor epegRNA R5. Alignments are filtered for reads overlapping the reporter locus. Linked alignments are excluded. **d.** As in c, but showing only linked alignments originating from reads without continuous alignment to the reference genome.

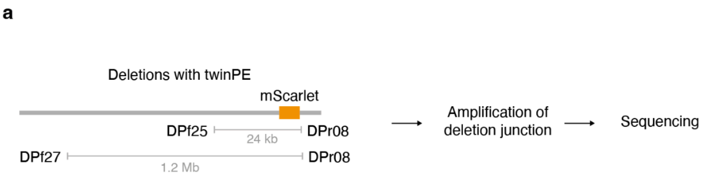


**b**

Deletion outcome

| Reads | intended edit | mutated attB site | other outcomes |
|---|---|---|---|
| | DPf25 + DPr08 | | |
| | 383,713 (96.4%) | 8,705 (2.2%) | 5,738 (1.4%) |
| | DPf27 + DPr08 | | |
| | 462,140 (94.6%) | 17,728 (3.6%) | 8,737 (1.8%) |

**Figure S4: Editing outcomes at the twinPE deletion junction. a.** Schematic of generated deletions. **b.** Read counts and frequencies for the different editing outcomes.

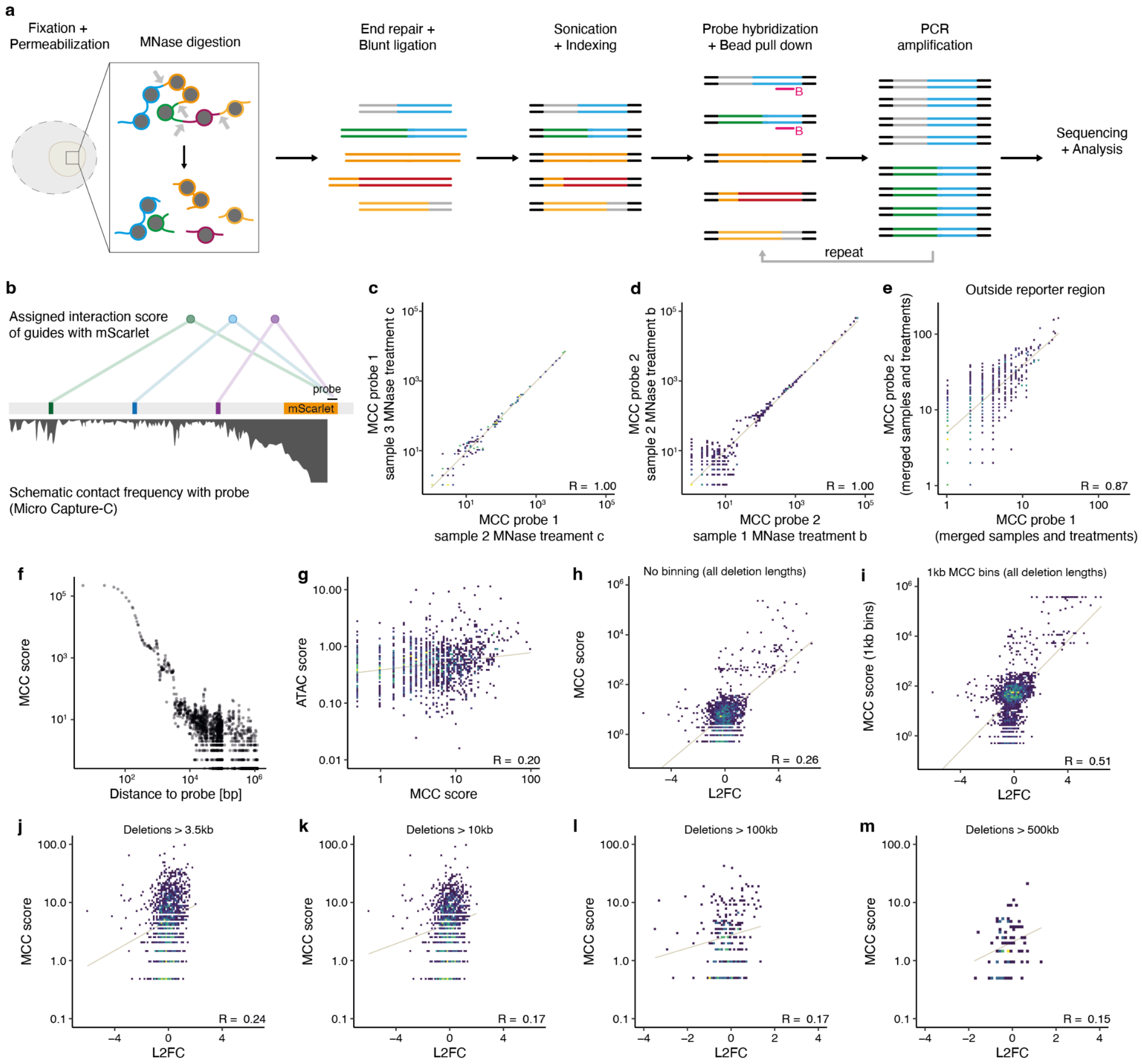


**Figure S5: Micro-capture C workflow, reproducibility, and correlation with screen features. a.** Schematic of the Micro Capture-C protocol (Hamley et al. 2023). The cells are fixed and permeabilized to digest genomic DNA with micrococcal nuclease (MNase). The fragments are end-repaired and ligated, sonicated, and indexed for sequencing. Biotinylated probes (B) are hybridized to the target region, and pull-down is performed with streptavidin beads. The enriched sample is amplified with PCR, and enrichment is repeated before sequencing and analysis. **b.** Schematic of MCC score assignment to the guides based on contact frequency with the probe hybridized to the reporter region. **c.** Scatter plot of MCC scores for replicates with probe 1. MNase treatment c: 15 Kunitz Units. R: Pearson's R. **d.** As (c) but for probe 2. MNase treatment b: 10 Kunitz Units. **e.** Correlation between probe 1 and probe 2 interaction scores for merged MNase treatments and samples for target sites >3.5kb from the probe region. **f.** Contact frequency score (MCC score, y-axis) compared to the distance to the probe (x-axis). **g.** Correlation between MCC score (x-axis) and ATAC signal R: Pearson's R. **h,i.** Correlation between MCC score (y-axis) and editing rates as log2 fold change (L2FC, x-axis) for different binning strategies (h: no binning, i: 1000 bp bins). R: Pearson's R. **j-m.** As e, but for different deletion length thresholds: 3.5kb (j), 10kb (k), 100kb (l), and 500kb (m).

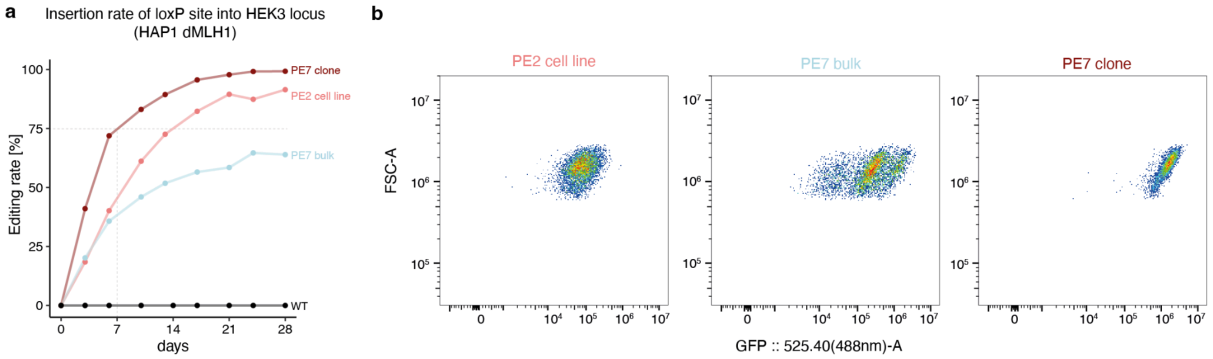


**Figure S6: HAP1-PE7 cell line generation. a.** Time course in days (x-axis) measuring the insertion rate (y-axis) of the 34nt loxPsym site into the HEK3 genomic target site for the HAP1-PE7 clone (dark red), bulk HAP1-PE7 (blue), HAP1 wild-type (WT, black), and HAP1-PE2 (PE2, light red). **b.** Flow cytometry measurement of GFP levels (x-axis) and forward scatter (FSC-A, y-axis) on day 28 for PE2 (light red, left panel), PE7 bulk (blue, middle panel), and HAP1-PE7 clone aD8 (dark red, right panel).

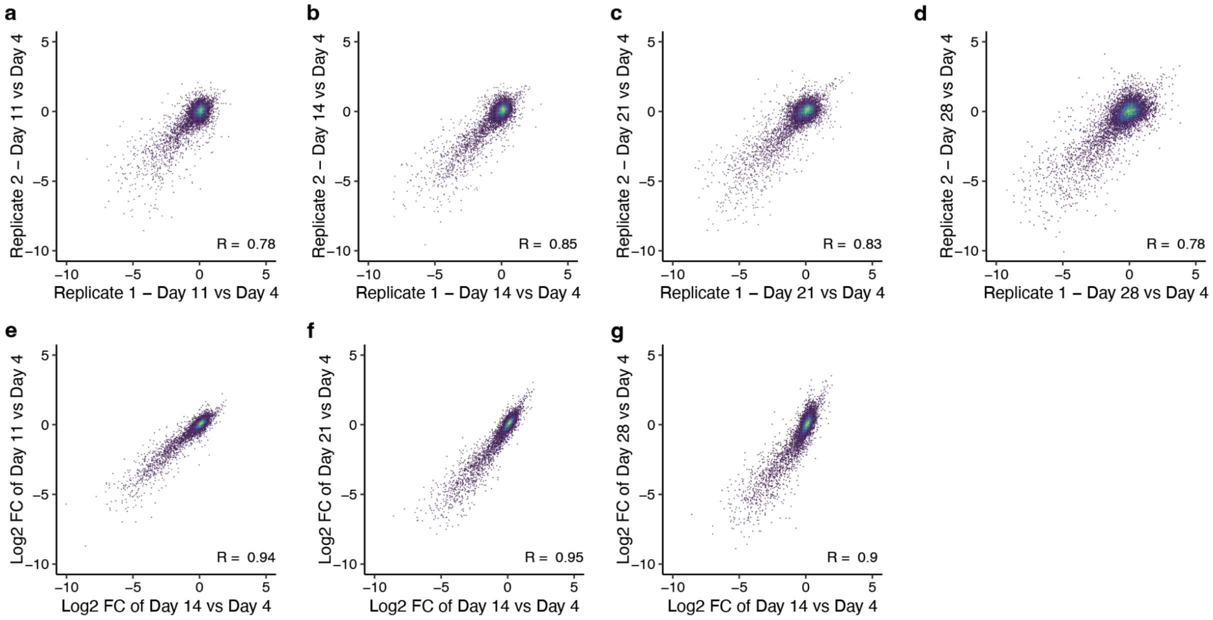


**Figure S7: Correlation of replicates and time points in the paired epegRNA screen. a-d.** Scatterplot showing the correlation between replicate 1 (x-axis) and replicate 2 (y-axis) for log2 fold changes (LFC) on day 11, day 14, day 21, and day 28 compared to day 4. R: Pearson's R. **e-g.** Correlation between merged replicates for LFC on day 11, day 21, and day 28 compared to day 4 (y-axis) and day 14 compared to day 4 (x-axis). R: Pearson's R.

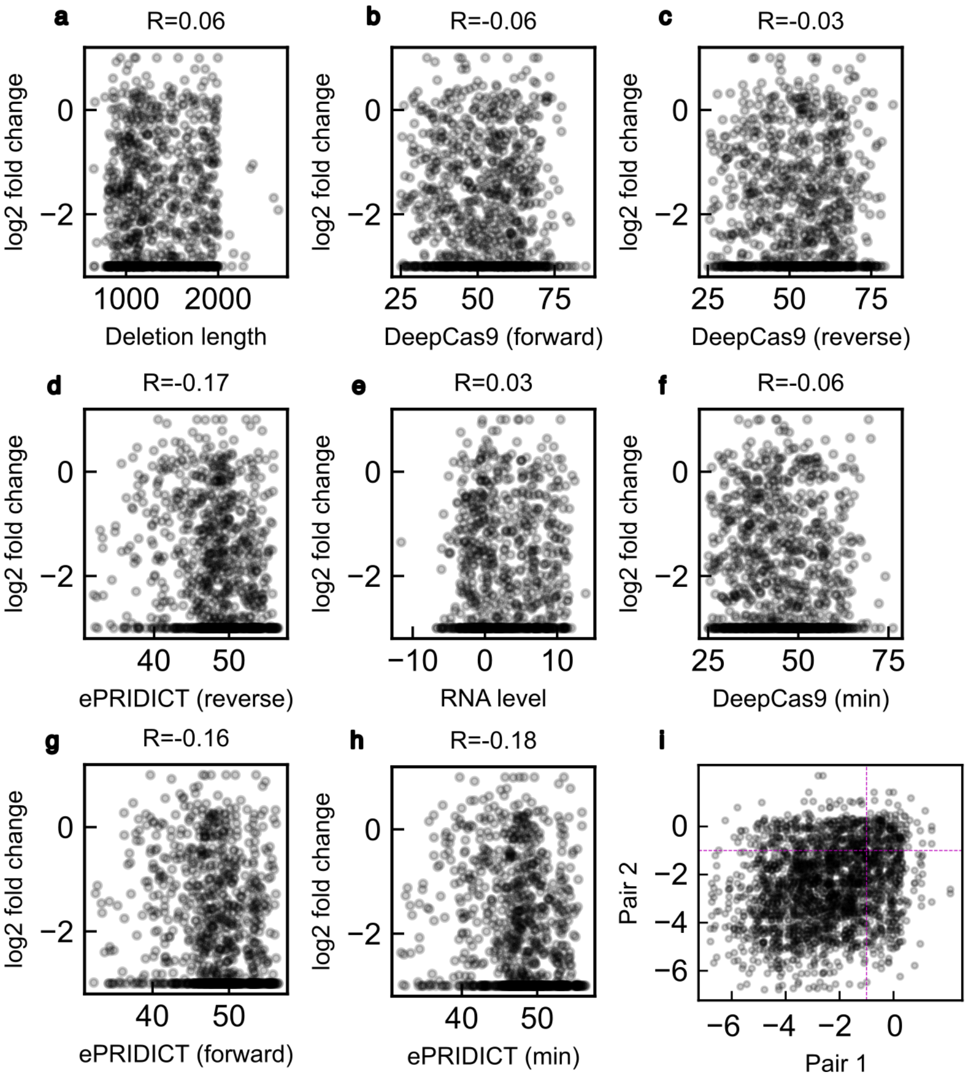


**Figure S8: Correlates of log-fold changes in the paired epegRNA screen.**

**a-h.** Log-fold change in the screen (y-axis; censored to (-3,1)) contrasted against a score for the pair of epegRNAs used to generate it (x-axis) for different deletions (markers. **i.** Log-fold change in the screen (x- and y-axis) from pairs of deletions (markers) that overlap, and target essential gene (CRISPR screen LFC < -1) exons. Dashed lines: guides for value -1.

### Supplementary Tables

Table S1. pegRNA library used for reporter screen and their log-fold changes (separate file)
Table S2. Sequences of single pegRNAs and other constructs used
Table S3. Deletion model features
Table S4. Paired pegRNA library used for genome-wide screen (separate file).
Table S5. Sanger sequencing results for non-essential prime deletions
Table S6. Primers used
Table S7. Plasmids used

**Table S2**. Sequences of single pegRNAs and other constructs used

| Name | Spacer | Extension |
|---|---|---|
| *R1* | GCTTCAGCACGCCGTCCTCG | CAGCACGCCGTCCATGATCCTGACGACGGAGGTCGCCGTCGTC GACA |
| *R2* | GTACAGCTCGTCCATGCCGC | AGCTCGTCCATGCATGATCCTGACGACGGAGGTCGCCGTCGTC GACA |
| *R3* | GTGAGCTCGAGATCTGAGTC | GCTCGAGATCTGAATGATCCTGACGACGGAGGTCGCCGTCGTC GACA |
| *R4* | GCCGCTTCCTCTAGATCCGG | CTTCCTCTAGATCATGATCCTGACGACGGAGGTCGCCGTCGTC GACA |
| *F1* | GTTCCCCGAGGGCTTCAAGT | CGAGGGCTTCAGGCTTGTCGACGACGGCGGTCTCCGTCGTCAG GA |
| *F2* | GAGAAGCCCGTGCAGATGCC | CCCGTGCAGATGGCTTGTCGACGACGGCGGTCTCCGTCGTCAG GA |
| *NT* | GAACAAGATGGATTGCACGC | GATGGATTGCAGGCTTGTCGACGACGGCGGTCTCCGTCGTCAG GA |
| *F3* | GTAATCGTGCGAGAGGGCGC | GTGCGAGAGGGGGCTTGTCGACGACGGCGGTCTCCGTCGTCAG GA |
| *F6* | GATGGGAAGGCTCCTCTCCC | TCCTGACGACGGAGACCGCCGTCGTCGACAAGCCAGAGGAGCC TTCC |
| *F7* | GAACTTAATGTACTCATCTT | TCCTGACGACGGAGACCGCCGTCGTCGACAAGCCATGAGTACA TTAA |
| *F4* | GCTAGGCGAGGTTGCGGGAA | TCCTGACGACGGAGACCGCCGTCGTCGACAAGCCCCGCAACCT CGCC |
| *F5* | GTTTGTCGTCCTCCCCTCAC | TCCTGACGACGGAGACCGCCGTCGTCGACAAGCCAGGGGAGGA CGAC |

**Table S3**. Deletion model features

| **Name** | **Category** | **Description** |
|---|---|---|
| *Deletion length* | deletion | Length of the deletion |
| *MCC 1kb bin* | deletion | MCC score binned by 1000 bp |
| *percA_spc* | pegRNA | Percentage of A in the spacer |
| *percC_spc* | pegRNA | Percentage of C in the spacer |
| *percT_spc* | pegRNA | Percentage of T in the spacer |
| *percG_spc* | pegRNA | Percentage of G in the spacer |
| *percGC_ext* | pegRNA | Percentage of GC in the extension |
| *percGC_pbs* | pegRNA | Percentage of GC in the PBS |
| *percGC_spc* | pegRNA | Percentage of GC in the spacer |
| *VF_spc* | pegRNA | Predicted folding free energy of the spacer (VFold model) |
| *VF_ext* | pegRNA | Predicted folding free energy of the extension (VFold model) |
| *PRIDICT2* | prediction | PRIDICT2 score (HEK293T) for attB site insertion |
| *ePRIDICT* | prediction | ePRIDICT score (full model, K562) for target site |
| *DeepPrime* | prediction | DeepPrime score (HEK293T) for 3nt insertion |
| *DeepCas9* | prediction | DeepCas9 score for spacer |

| | | |
|---|---|---|
| *DeepSpCas9* | prediction | DeepSpCas9 score for spacer |
| *Doench2014OnTarget* | prediction | OnTarget score for spacer |
| *DoenchCFD* | prediction | Specificity score for spacer |
| *DNase (HEK293T)* | chromatin | ENCFF969MBJ |
| *ATAC (HAP1)* | chromatin | GSE111047 |
| *H3K27me3 (HEK293T)* | chromatin | SRR8937480 |
| *H3K27me3 (HAP1)* | chromatin | ENCFF898ETI |
| *H3K9me3 (HEK293T)* | chromatin | SRR11453034 |
| *H3K9me3 (HAP1)* | chromatin | ENCFF782DHX |

**Table S5**. Sanger sequencing results for non-essential prime deletions

| Deletion name | Reference with alignment |
|---|---|
| *BTF3_1* | https://benchling.com/s/seq-OUGOBQvAcKEXa8QSA4bz?m=slm-6xBeUzgM5yrWgx4qap6W |
| *BTF3_2* | https://benchling.com/s/seq-OUGOBQvAcKEXa8QSA4bz?m=slm-6xBeUzgM5yrWgx4qap6W |
| *BTF3_3* | https://benchling.com/s/seq-OUGOBQvAcKEXa8QSA4bz?m=slm-6xBeUzgM5yrWgx4qap6W |
| *DDB1_1* | https://benchling.com/s/seq-MdBhyI3NjzuAgtfi1gvz?m=slm-Q18WkAn3HYA20pB8nZcx |
| *DDB1_2* | https://benchling.com/s/seq-MdBhyI3NjzuAgtfi1gvz?m=slm-Q18WkAn3HYA20pB8nZcx |
| *EIF4B_1* | https://benchling.com/s/seq-q7N5gj5O3ihcOmFnfxPq?m=slm-DsbdHkCwk7BZKQsoPXwe |
| *EIF4B_2* | https://benchling.com/s/seq-QdOn7Tz85qEvDz7g9dOI?m=slm-S3Is1u2s5PippqtsIK0b |
| *EIF4G1* | https://benchling.com/s/seq-sCjktEfJ20YrUGFupNCi?m=slm-EVOmi17kVYn01CK5R9sf |
| *IntChr11* | https://benchling.com/s/seq-4ydDa5Eo1WgEzwU6GDPU?m=slm-MPd1dBxXTxq71UbZm2Jp |
| *KPNB1_1* | https://benchling.com/s/seq-qrMy6CVzKaMCgS3azBLd?m=slm-8xR3Ph6OtIpRojPS8PtI |
| *KPNB1_2* | https://benchling.com/s/seq-qrMy6CVzKaMCgS3azBLd?m=slm-8xR3Ph6OtIpRojPS8PtI |
| *KPNB1_3* | https://benchling.com/s/seq-qrMy6CVzKaMCgS3azBLd?m=slm-8xR3Ph6OtIpRojPS8PtI |
| *NC_1* | https://benchling.com/s/seq-jBUKezoQgHiRypsYvEnp?m=slm-5F704AxLhvWdFJZja6re |
| *NC_2* | https://benchling.com/s/seq-rS183hJNwDkelRat0sbr?m=slm-JoJUryaoAdhCLEhwFCk4 |
| *PMAIP1_1* | https://benchling.com/s/seq-k2yKrB8G4fR0xfMd1FVD?m=slm-MpAS61pQEjOtkBM1g9nR |
| *PMAIP1_2* | https://benchling.com/s/seq-k2yKrB8G4fR0xfMd1FVD?m=slm-MpAS61pQEjOtkBM1g9nR |
| *SRSF1_1* | https://benchling.com/s/seq-ekcsql6m7a6p8vlfV0zj?m=slm-3WuGB4GkEkLTQ7JDvria |
| *SRSF1_2* | https://benchling.com/s/seq-ekcsql6m7a6p8vlfV0zj?m=slm-3WuGB4GkEkLTQ7JDvria |
| *TXNRD1_1* | https://benchling.com/s/seq-DJbf3tRVvJjMAvCuJYxq?m=slm-QGHhDn1Fzep39HvNYfNS |
| *TXNRD1_2* | https://benchling.com/s/seq-DJbf3tRVvJjMAvCuJYxq?m=slm-QGHhDn1Fzep39HvNYfNS |

| | |
|---|---|
| *TXNRD1_3* | https://benchling.com/s/seq-DJbf3tRVvJjMAvCuJYxq?m=slm-QGHhDn1Fzep39HvNYfNS |

**Table S6**. Primers used

*Homology arms for forward-strand targeting pegRNA in library*

| HA name | Sequence |
|---|---|
| *attB_PASTE* | GGCCGGCTTGTCGACGACGGCGGTCTCCGTCGTCAGGA |
| *attB_34* | GGCTTGTCGACGACGGCGGTCTCCGTCGTCAGGA |
| *attB_29* | GGCTTGTCGACGACGGCGGTCTCCGTCGT |
| *attB_mod1* | CGCCCGCCGACGACGGCGGTCTCCGTCGT |
| *attB_mod2* | GACTTATCTACGACGGCGGTCTCCGTCGT |

*pegRNA scaffolds*

| Scaffold | Sequence |
|---|---|
| *Improve* | GTTTAAGAGCTATGCTGGAAACAGCATAGCAAGTTTAAATAAGGCTAGTCCGTTATCAACTTGAAAAAGTGGCACCGAGTCGGTGC |
| *PASTE* | GTTTGAGAGCTATGCTGGAAACAGCATAGCAAGTTCAAATAAGGCTAGTCCGTTATCAACTTGAAAAAGTGGCACCGAGTCGGTGC |

*Primer list*

| Primer ID | Sequence |
|---|---|
| *962* | ACACTCTTTCCCTACACGACGCTCTTCCGATCTGATGGCTTTATATATCTTGTGGAAAGGACGAAACACC |
| *963* | ACACTCTTTCCCTACACGACGCTCTTCCGATCTCTAGAATGGCTTTATATATCTTGTGGAAAGGACGAAACACC |
| *1396* | GTGACTGGAGTTCAGACGTCTGCTCTTCCGATCTTGACCGCTGAAGTACAAGTGGT |
| *1101* | ACCAAGGAAAGTCGATGGCGTA |
| *1107* | GTAAGAACAAGCGGACGTTGGG |
| *884* | CCCAGCCAAACTTGTCAACC |
| *1752* | AACCACCGCGACCTCAGTGGTG |
| *1753* | GACCAGACAAACCGGG |
| *860* | ACACTCTTTCCCTACACGACGCTCTTCCGATCTGGCTTTATATATCTTGTGGAAAGGACGAAACACC |
| *964* | ACACTCTTTCCCTACACGACGCTCTTCCGATCTTGGACACATGGCTTTATATATCTTGTGGAAAGGACGAAACACC |
| *883* | ATGTGGGCTGCCTAGAAAGG |

*Cas9 guides for targeted enrichment with nanopore sequencing*

| Guide ID | Target sequence |
|---|---|

| | |
|---|---|
| *Cas9guide_Dipper_rev_2* | TTTAAGATCCTGGCAGCAAGAGG |
| *Cas9guide_Dipper_rev_3* | AGAGCTCTATTGGAAATCAGTGG |
| *Cas9guide_Dipper_for_2* | TGCGATCTGCTGTCTGCAAACGG |
| *Cas9guide_Dipper_for_3* | GCACTGGTTAGATTTGGCAGAGG |

Gene Fragments

| Name | Sequence |
|---|---|
| *GF1* | gtaagccggtccaccaacaaagcgCGTCTCCGCGGTTCTATCTAGTTACGCGTTAAACCAACTAGAATTTTTTTTTCTAGAGATCCGACGCCGCCATCTCTAGGCCCGCGCCGGCCCCCTCGCACAGACTTGTGGGAGAAGCTCGGCTACTCCCCTGCCCCGGTTAATTTGCATATAATATTTCCTAGTAACTATAGAGGCTTAATGTGCGATAAAAGACAGATAATCTGTTCTTTTTAATACTAGCTACATTTTACATGATAGGCTTGGATTTCTATAAGAGATACAAATACTAAATTATTATTTTAAAAAACAGCACAAAAGGAAACTCACCCTAACTGTAAAGTAATTGTGTGTTTTGAGACTATAAATATGCATGCGAGAAAAGCCTTGtGAGACGgcggttgtcattgaaggagctgcc |

MCC probe sequences

| MCC probe | Sequence |
|---|---|
| *probe_mS_1* | TCCGAGGGCCGCCACTCCACCGGCGGCATGGACGAGCTGTACAAGTCCGGACTCAGATCTCGAGCTCAAGCTTCGAATTCTGCAGTCGACGGTACCGCGGGCCCGGGATCCACCGGATCT |
| *probe_mS_2* | GACATCACCTCCCACAACGAGGACTACACCGTGGTGGAACAGTACGAACGCTCCGAGGGCCGCCACTCCACCGGCGGCATGGACGAGCTGTACAAGTCCGGACTCAGATCTCGAGCTCAA |

### **Table S7**. Plasmids used

| Plasmid name | Description | Benchling link |
|---|---|---|
| *pCLYBL_CAGG_mScarlet* | Generating mScarlet cell line | https://benchling.com/s/seq-5QJfAd7y9D7XA6A0D9xf?m=slm-qdVazdUd4gL7sXV1nL1v |
| *pCLYBL_CAGG_mScarlet-degron* | Generating mScarlet-degron cell line | https://benchling.com/s/seq-cP21bnwHZuLxwzPLC25a?m=slm-yALiMvsV8o9KniWV1EkU |
| *pCAG-Cas9-U6-CLYBLgRNA* | Cas9 targeting the CLYBL locus | https://benchling.com/s/seq-s1oK76uG7RRENWtNtFxT?m=slm-rRLDhWN8CU3DylLkhWmt |
| *pLentiGuide-BlastR* | Lentiviral backbone for pegRNA with blasticidin resistance | https://benchling.com/s/seq-uiF1iq3pcEUBjZHtbIyw?m=slm-nNXNR7rpFJKjDCSTXfxm |
| *pU6-pegRNA-GG-acceptor* | Backbone for pegRNA with puromycin resistance | https://benchling.com/s/seq-HDuKt4H8pIVv3ytBcDrl?m=slm-sRGRa57a1b8nypxcxAPr |

| | | |
|---|---|---|
| *pU6-epegRNA-GG-acceptor* | Backbone for epegRNA | https://benchling.com/s/seq-flUbGgSl2GCGqoiiHgGd?m=slm-lV9fnvKFic7mnnJ56ago |
| *pLenti_epeg_acceptor* | Lentiviral backbone for epegRNA with puromycin resistance | https://benchling.com/s/seq-XGTdD2V78LDXmLQAoxhP?m=slm-52yiQ33DwOlTmLEH6yQK |
| *pLenti_PE7-GFP* | *Lentiviral plasmid with PE7 under SFFV promoter for generating the PE7 cell line* | *https://benchling.com/s/seq-14hRjHnvueDnd6EVSAW1?m=slm-E7CFScjY5gicbsGlwGZu* |
| *pLenti-puro-mCherry-epeg* | Backbone for epegRNA with puromycin resistance and mCherry | https://benchling.com/s/seq-Pa4toV336iUEMXnyIFH7?m=slm-MmiuZaF5U42qziWGH6B0 |
| *Hygro-BFP-attP (GT)* | Donor plasmid with attP site | https://benchling.com/s/seq-GEejZpHo6JNlINMNgUFN?m=slm-1FWBWogtGHyssDhXIncM |
| *evoBxb1* | Plasmid for expressing evolved Bxb1 in the mammalian cells | Addgene: 222338 |
| *wtBxb1* | Plasmid for expressing wild-type Bxb1 in the mammalian cells | Addgene: 222337 |
| *eeBxb1* | Plasmid for expressing engineered Bxb1 in the mammalian cells | Addgene: 222339 |
| *wtBxb1 (Parts Lab)* | Plasmid for expressing wild-type Bxb1 in the mammalian cells | Addgene: 51552 |
| *pLenti-puro-mCherry* | Lentiviral backbone with puromycin resistance and mCherry | https://benchling.com/s/seq-iAU0Hnrpg4LEMuTMiqep?m=slm-KgKbzKeJdyBBrU5PI0ax |
